# Compression of quantification uncertainty for scRNA-seq counts

**DOI:** 10.1101/2020.07.06.189639

**Authors:** Scott Van Buren, Hirak Sarkar, Avi Srivastava, Naim U. Rashid, Rob Patro, Michael I. Love

**Affiliations:** University of North Carolina-Chapel Hill; University of Maryland; New York Genome Center

## Abstract

**Motivation:** Quantification estimates of gene expression from single-cell RNA-seq (scRNA-seq) data have inherent uncertainty due to reads that map to multiple genes. Many existing scRNA-seq quantification pipelines ignore multi-mapping reads and therefore underestimate expected read counts for many genes. *alevin* accounts for multi-mapping reads and allows for the generation of “inferential replicates”, which reflect quantification uncertainty. Previous methods have shown improved performance when incorporating these replicates into statistical analyses, but storage and use of these replicates increases computation time and memory requirements.

**Results:** We demonstrate that storing only the mean and variance from a set of inferential replicates (“compression”) is sufficient to capture gene-level quantification uncertainty. Using these values, we generate “pseudo-inferential” replicates from a negative binomial distribution and propose a general procedure for incorporating these replicates into a proposed statistical testing framework. We show reduced false positives when applying this procedure to trajectory-based differential expression analyses. We additionally extend the *Swish* method to incorporate pseudo-inferential replicates and demonstrate improvements in computation time and memory consumption without any loss in performance. Lastly, we show that the removal of multi-mapping reads can result in significant underestimation of counts for functionally important genes in a real dataset.

**Availability and implementation:** *makeInfReps* and *splitSwish* are implemented in the development branch of the R/Bioconductor *fishpond* package available at http://bioconductor.org/packages/devel/bioc/html/fishpond.html. Sample code to calculate the uncertainty-aware *p*-values can be found on GitHub at https://github.com/skvanburen/scUncertaintyPaperCode.

## 1 Introduction

Single-cell RNA-sequencing (scRNA-seq) allows for analysis of gene expression data at the level of individual cells. This cell-level expression is often summarized in terms of expected read counts for each gene. Many scientific questions that were previously difficult to address using bulk RNA-seq can now be directly studied with scRNA-seq, including direct identification of complex and rare cell populations as well as the analysis of cellular development trajectories (Hwang *et al*., 2018). However, common pipelines for obtaining gene-level expression estimates for scRNA-seq either discard multi-mapping reads entirely, which may lead to biased quantification estimates, or have no means to evaluate the quantification uncertainty in expression estimates that is imparted by such reads (Srivastava *et al*., 2019).

*alevin* (Srivastava *et al*., 2019) is a droplet-based scRNA-seq (dscRNA-seq) quantification pipeline that builds upon *Salmon* (Patro *et al*., 2017) and improves upon prior pipelines for scRNA-seq in several important ways. First, *alevin* is able to quantify reads that map to multiple genes by first resolving multi-mapping reads using a parsimony criterion, and then resolving equally-parsimonious solutions by use of the EM algorithm. This reduces systematic biases in quantified gene-level counts (Srivastava *et al*., 2019). Compared to paired-end bulk RNA-seq, dscRNA-seq exhibits a 3’ coverage bias and generates one read of transcript sequence, which worsens the impact of multi-mapping reads relative to typical bulk experiments. Combined, these effects may result in as many as 23% of reads mapping to multiple genes, which are discarded by default by alternate quantification methods that do not employ the EM algorithm, such as *dropEst* (Petukhov *et al*., 2018), *Cell Ranger* (Zheng *et al*., 2017), *STARsolo* (Dobin *et al*., 2013), or *bustools* (Melsted *et al*., 2019). When compared to existing scRNA-seq quantification pipelines, *alevin* improved the accuracy of quantification results when comparing pseudo-bulk samples of mouse retina data generated with scRNA-seq to bulk RNA-seq of the same tissue type (Srivastava *et al*., 2019). Improvement was greatest for genes with lower levels of sequence uniqueness (higher potential for multi-mapping reads), and lower for genes with 100% uniqueness (lowest potential for multi-mapping reads). We demonstrate later that discarding multi-mapping reads can result in significant underestimation of counts for functionally important genes using recently published scRNA-seq data of developing mice embryos (Pijuan-Sala *et al*., 2019).

Second, *alevin* can additionally assess the inherent quantification uncertainty in cell-level expected read counts caused by multi-mapping reads by examining the distribution of quantification estimates derived from bootstrap replicates from the original set of reads. Bayesian models for expression estimates alternatively may draw replicates directly from a corresponding posterior distribution, often using MCMC methods such as Gibbs sampling. These two types of replicates can be collectively referred to as “inferential replicates,” and either type provides a relative measure of the level of quantification uncertainty. Inferential replicates have been previously used in bulk RNA-seq to capture inferential uncertainty of gene or transcript-level quantification estimates (Turro *et al*., 2011; Li and Dewey, 2011; Al Seesi *et al*., 2014; Bray *et al*., 2016; Mandric *et al*., 2017; Pimentel *et al*., 2017; Patro *et al*., 2017; Froussios *et al*., 2019; Zhu *et al*., 2019; Tiberi and Robinson, 2020; Van Buren and Rashid, 2020). By default, *alevin* stores only the sample mean and variance of the bootstrap replicates for each gene and cell instead of the full set of replicates. This “compression” procedure greatly reduces the amount of disk space and memory required for storage and down-stream analysis. However, it has not been evaluated whether this procedure sufficiently captures the quantification uncertainty reflected in a full set of inferential replicates, thereby justifying the avoidance of their storage and direct use in downstream analyses.

In this paper we demonstrate that storage of only the mean and variance of the bootstrap replicates is sufficient to capture the gene-level inferential uncertainty, greatly reducing the amount of disk space, memory, and load time required for downstream analyses. We additionally extend the *Swish* method to operate on “pseudo-inferential” replicates drawn from a negative binomial distribution using stored compression parameters. We show that the use of pseudo-inferential replicates has comparable performance to results that instead utilized bootstrap replicates. Lastly, we evaluate the impact of accounting for quantification uncertainty into trajectory-based scRNAseq differential expression analysis using *tradeSeq* (Van den Berge *et al*., 2020), and demonstrate that improvements in the false discovery rate (FDR) can be obtained by incorporating pseudo-inferential replicates.

## 2 Methods

### 2.1 Uncertainty aware scRNA-seq workflow with compression

A summary of the uncertainty-aware scRNA-seq workflow with compression is given in Figure 1. A list of FASTQ files originating from a dscRNA-seq experiment are utilized as input, where *alevin* is run with the flag `--numCellBootstraps 20` to conduct the quantification and store the mean and variance of 20 bootstrap replicates from each gene and cell. Under this setting, the bootstrap replicates are not retained. We additionally evaluated the use of 100 bootstrap replicates instead of 20. Parameters for a negative binomial distribution are then derived from these compressed estimates (see Section 2.3 for more detail) and are used to sample pseudo-inferential replicates for use in various downstream tasks in lieu of the original bootstrap replicates. Pseudo-inferential replicates can be generated separately for each gene, allowing tasks such as differential expression analysis to be easily distributed across separate CPUs or jobs.

**Figure 1:**
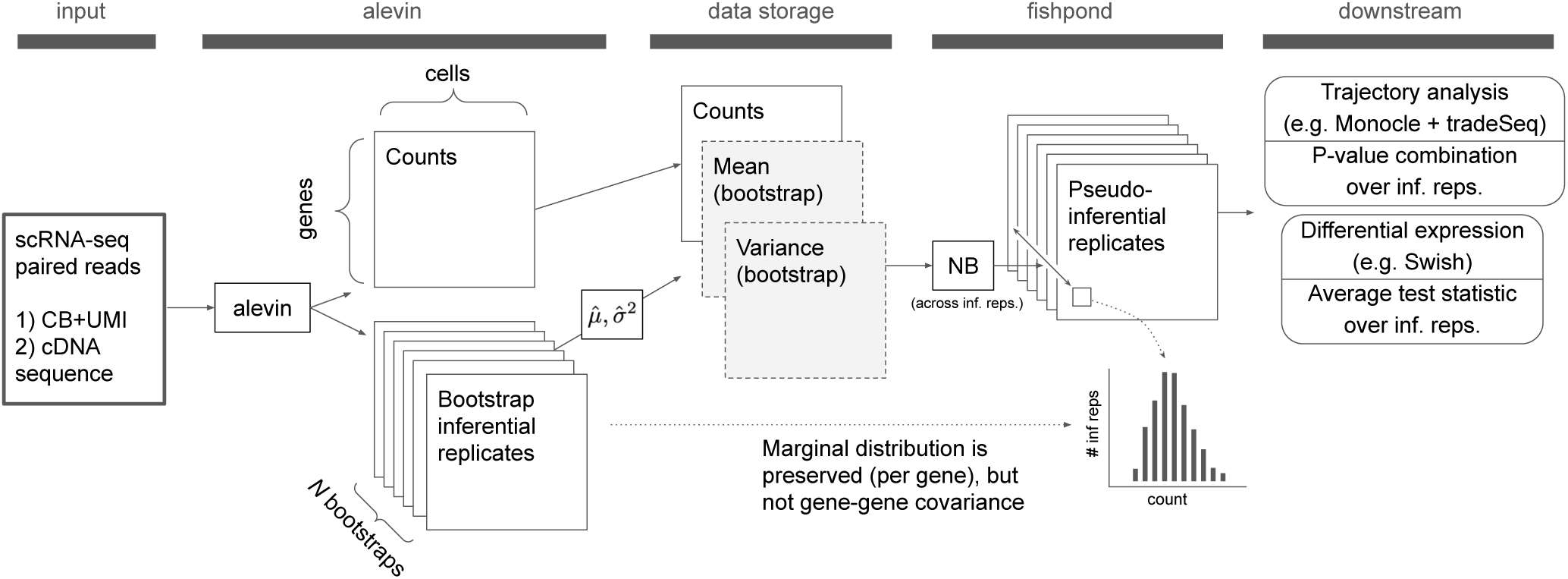
Compression of scRNA-seq quantification uncertainty. This procedure stores solely the mean and variance of the bootstrap replicate count matrices, with this compressed information later used to regenerate marginal (per-gene) pseudo-inferential replicates as needed. (Abbreviations: CB - cell barcode, UMI - unique molecular identifier, NB -negative binomial.)

### 2.2 Simulation procedure

Using statistical simulation, we evaluated the performance benefit of using compression to incorporate quantification uncertainty into standard group-based scRNA-seq differential expression analysis. We also evaluated the performance benefit for trajectory-based differential expression analysis. Trajectory analysis for scRNA-seq is an important development that enables study of the collection of paths, or lineages, in which a cell of one type differentiates into a new cell type (Cannoodt *et al*., 2016; Saelens *et al*., 2019). A collection of lineages is often referred to as a “trajectory,” and many methods are available to conduct trajectory-based differential expression analysis, including *tradeSeq* (Van den Berge *et al*., 2020). *tradeSeq* fits a separate modified generalized additive model (GAM) (Hastie and Tibshirani, 1986) to expression values for each gene to model how values change across lineages and “pseudotimes,” temporal variables that are not measured in exact units but index movement from the beginning of a lineage towards the end. Here, *tradeSeq* was used in combination with pseudo-inferential replicates such that the analysis would be sensitive to quantification uncertainty. See Section 2.4 for implementation specifics.

To simulate data under simple two-group differences for the former scenario, we utilized the *Splat* method from the *splatter* package (Zappia *et al*., 2017). Similar to Zhu *et al*. (2019), we set the DE *factor location* parameter to be 3 on the log_2_ scale, the DE *factor scale* parameter to be 1 on the log_2_ scale, 10% of genes to be differentially expressed, and simulated data for 100 cells in each of two groups. To simulate data under the latter trajectory-based scenario, we used the *dynverse* framework that was previously used to benchmark trajectory inference methods (Saelens *et al*., 2019), and was also used in benchmarking *tradeSeq* (Van den Berge *et al*., 2020). In particular, we considered “bifurcating” and “trifurcating” trajectories, similarly to Van den Berge *et al*. (2020), for both 100 and 250 cells. We set the level of differential expression to be 20%, as in the *tradeSeq* paper. Both simulations used 60,179 genes, corresponding to the number of genes from the GENCODE version 32 annotation from the reference chromosomes only (Harrow *et al*., 2012; Frankish *et al*., 2018) that were able to be quantified by *alevin*. Simulated counts were assigned to actual genes based on the rank of the gene’s average expression from quantification of a dataset of peripheral blood mononuclear cells (PBMC), specifically the publicly available PBMC 4k dataset (Zheng *et al*., 2017). This procedure preserved the rank of the genes by expression across simulated and real data.

Following the generation of gene-level counts, we utilized the *minnow* framework (Sarkar *et al*., 2019) to simulate realistic scRNA-seq reads corresponding to the simulated counts from *splatter* or *dynverse. minnow* is able to simulate dscRNA-seq reads accounting for important characteristics of real dscRNA-seq data, including polymerase chain reaction amplification, cellular barcodes (CBs) and CB errors, unique molecular identifiers (UMI) for each read, and sequence fragmentation. *minnow* importantly is able to account for realistic patterns of uncertainty and multi-mapping of reads by its use of a (compact) De Bruijn graph instead of sampling reads directly from transcript sequences. The rates of multi-mapping used in sampling sequences from the De Bruijn graph were estimated from the aforementioned PBMC 4K dataset. The resulting dscRNA-seq reads were then quantified with *alevin*, and 20 bootstrap replicates of gene expression values were generated for each cell. We additionally evaluated the use of 100 bootstrap replicates instead of 20. All results utilized annotation files corresponding to the previously discussed annotation corresponding to the reference chromosomes only from GENCODE version 32. Quantified data for the trajectory analysis simulations were imported into *R* using the *tximport* package (Soneson *et al*., 2016) to obtain simple list output and data from the simple two group difference simulation were imported using the *tximeta* package (Love *et al*., 2020) to obtain *SummarizedExperiment* objects to simplify use with the *Swish* method (Zhu *et al*., 2019).

### 2.3 Evaluation of bootstrap replicates

We compared the bootstrap replicates from *alevin* to the true simulated counts, evaluating the coverage of various intervals constructed from the bootstrap replicates. To correct for differences in total count per cell due to reads not aligning, we scaled the simulated counts for each cell to have the same total mapped count as from *alevin* before evaluating interval coverage. Additionally, *minnow* is unable to generate reads for genes whose transcript sequences are shorter than the simulated read length (101). Our simulation had 3,068 such genes, and we removed these genes from consideration before calculating coverage.

We considered 95% intervals constructed using the full set of bootstrap replicates and using quantiles from a negative binomial distribution whose parameters were determined from the mean and variance of the bootstrap replicates. If the latter interval type provided similar results to the former type, compression of the bootstrap replicates could be performed without a loss of relevant information. Note that negative binomial was used here for the distribution of counts for one gene and one cell across bootstrap replicates, not across genes or across cells. Specifically, let *V*_*igj*_ be the count for cell *i* = 1, …, *n*, for gene *g* = 1, …, *G*, and in bootstrap replicate *j* = 1, …, 20. If we let *V*_*ig*_ = (*V*_*ig*1_, …, *V*_*ig*{20}_) be the entire vector of bootstrap values for cell *i* and gene *g*, we constructed the former interval type for sample *i* and gene *g* as (*q*_0.025_, *q*_0.975_), where *q*_0.025_ and *q*_0.975_ are 0.025 and 0.975 quantile values of *V*_*ig*_ respectively. Since the 0.025 and 0.975 quantiles are not defined exactly with 20 values, standard interpolation techniques are used to estimate these quantiles (Hyndman and Fan, 1996). The latter interval type was constructed using a negative binomial distribution with parameters *µ* and *ϕ* chosen such that E(*Y*) = *µ* and 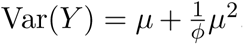. The parameter *ϕ* governs the amount of extra-Poisson dispersion, with large values of *ϕ* indicating a distribution closer to Poisson, and small values of *ϕ* associated with higher over-dispersion. Letting 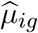 be the sample mean of *V*_*ig*_ and 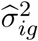 be the sample variance of *V*_*ig*_, we constructed the negative binomial-based interval for sample *i* and gene *g* as (*w*_0.025_, *w*_0.975_), where *w*_0.025_ and *w*_0.0975_ are the quantiles from a negative binomial distribution with 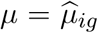 and 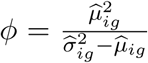. In practice, we set the maximum value of *ϕ* to be 1,000 when.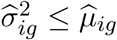.

The “coverage” for a given gene within a cell was defined as equal to one if the scaled, simulated count is contained in the interval and zero otherwise. The overall coverage for a gene was obtained by averaging the coverage values for the gene across all cells. In general, if the simulated replicates accurately reflected the true expression profile they were simulated from, we would expect coverage of the true count to be close to the nominal value, e.g. 95%. Additionally, if storage of only the mean and variance of the bootstrap replicates was sufficient to capture the gene-level inferential uncertainty present in the bootstrap replicates, then coverage of the two interval types should be similar. Both interval types are similar to Bayesian credible intervals (Hoff, 2009; Gelman *et al*., 2013), where the parameter of interest in our case would be the scaled, simulated count. However, note that the use of bootstrap replicates to construct the intervals means these intervals cannot be thought of as proper credible intervals since no posterior distribution is used in their construction. We only considered genes that had counts of at least 10 in at least 10 cells in our main coverage evaluations. This is because count values of zero proved substantially easier to cover than positive counts, as we will demonstrate later, resulting in very lowly expressed genes overly inflating coverage statistics when included.

To summarize the amount of quantification uncertainty present per cell and per gene, we utilized the inferential relative variance (InfRV) statistic proposed by Zhu *et al*. (2019). This quantity is defined for each cell and gene combination as:

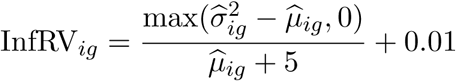

where 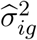 and 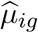 are the sample variance and sample mean values of the bootstrap replicates for cell *i* and gene *g* respectively. This quantity is roughly independent of the range of the counts, and the quantities 5 and 0.01 are respectively added to stabilize the statistic and ensure the final quantity is strictly positive for log transformation. The final InfRV value for a gene can then be taken as the average of each cell-specific value for the gene. The InfRV statistic is not directly incorporated into any testing procedure but instead is used to categorize genes based on quantification uncertainty for plotting and to evaluate how methods perform across differing levels of quantification uncertainty.

### 2.4 Incorporation of uncertainty into scRNA-seq trajectory analysis

Pseudo-inferential replicates were generated from a negative binomial distribution using distributional parameter values derived from the compressed uncertainty estimates, as detailed in Section 2.3. Lineages and pseudotimes were fit using the *slingshot* method (Street *et al*., 2018), and *trade-Seq* was used to fit the GAMs to expression counts utilizing these lineages and pseudotimes. The procedure was repeated on each replicate, and results were combined across replicates using two different approaches described in more detail below. We utilized the pre-defined `associationTest` and `patternTest` within the *tradeSeq* method to test for general differences in expression within a single lineage and between several distinct lineages respectively. We additionally utilized the `startVsEndTest` to test for differences in expression between the start and end of lineages and the `diffEndTest` to test for differences in expression between separate lineages near the end of the lineages. The fitting and testing procedure was repeated on 20 pseudo-inferential replicates simulated from a negative binomial distribution with parameters calculated according to the procedure discussed in Section 2.3. We considered several methods to combine results for a gene across the simulated datasets.

The first method was motivated from *Swish* (Zhu *et al*., 2019) in that it uses the mean test statistic over inferential replicates as its final test statistic. In contrast to *Swish*, which uses permutation to determine significance, the mean test statistic is compared to a parametric null distribution to determine significance. Specifically, *tradeSeq* utilizes Wald test statistics, which follow a chi-squared null distribution, for each of its significance tests. However, the associated degrees of freedom (df) of the chi-squared null distributions can change across genes and replicates for certain tests. To account for this, we first transformed *p*-values across replicates to a chi-squared distribution with df equal to the most commonly observed df value over the pseudo-inferential replicates. While the mean of chi-squared random variables do not follow a chi-squared distribution, we assumed the mean test statistic across replicates corresponds to a single hypothesis test for the gene of interest. We then were able to compare this mean test statistic to the same chi-squared distribution used in the inverse *p*-value transformation above to calculate final *p*-values for each gene determine significance. Note that the final *p*-values will not necessarily follow a uniform distribution under the null hypothesis with this approach. This method is referred to in Results as “MeanStatAfterInvChiSq.”

The second approach selects a specific percentile of the vector of raw *p*-values across replicates to be the final *p*-value for each gene and performs FDR correction on these selected *p*-values to determine significance. We considered the 50th and 75th percentiles, and refer to these methods in Results as “Pval50Perc” and “Pval75Perc” respectively. This procedure is similar to the procedure utilized by *RATs* (Froussios *et al*., 2019), which tests for differential transcript usage (DTU) in bulk RNA-seq data. *RATs* incorporates inferential uncertainty by requiring a certain proportion (default 0.95) of FDR-adjusted *p*-values across inferential replicates (either Gibbs or bootstrap) to show significance at a given nominal FDR level for the gene to be considered to show significant DTU. However, this approach requires the full set of FDR-adjusted *p*-values across inferential replicates to be retained if significance is to be evaluated at a different FDR threshold. Depending on the number of significance tests and inferential replicates used, the disk space and memory required to store and load all *p*-values could be prohibitive. In contrast, our proposed approach enables evaluation of multiple FDR cutoffs while only requiring storage of a single *p*-value for each significance test. We will demonstrate later that our proposed approach provides very similar performance in practice to the one utilized by *RATs*.

### 2.5 Modification of *Swish* to use pseudo-inferential replicates

We additionally modified the existing *Swish* implementation (Zhu *et al*., 2019) to enable it to use pseudo-inferential replicates generated from a negative binomial distribution. This can greatly reduce the amount of disk space and memory required to incorporate inferential replicate information into existing analyses. Pseudo-inferential replicates can be simulated using the `makeInfReps` function in the `fishpond` Bioconductor package. The `splitSwish` function was also added to the package, and allows most of the *Swish* computations to be distributed across cores using *Snakemake* (Köster and Rahmann, 2012). Results from each core are gathered prior to calculation of the final *q*-value, using the `qvalue` package and function (Storey, 2002). Only the compressed inferential statistics 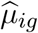 and 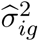 are sent to each core, with pseudo-inferential replicates generated and used as needed per core. This further reduces total memory and running time per job.

### 2.6 Simulation evaluation

To evaluate the performance of the previously discussed simulations, we used the *iCOBRA* package (Soneson and Robinson, 2016) to generate plots that compare the true positive rate (TPR) across different false discovery rates (FDR) at nominal FDR thresholds of 1%, 5%, and 10%. We additionally stratified the plots based on InfRV to compare performance across differing levels of quantification uncertainty.

### 2.7 Mouse embryo data

We evaluated the effect of multi-mapping reads and quantification uncertainty on trajectory-based differential expression with data from a recent scRNA-seq study by Pijuan-Sala *et al*. (2019). This study sequenced RNA from 116,312 single cells from mouse embryos, collected at nine sequential time points that range from 6.5 to 8.5 days post-fertilization. We considered data at a subset of time points, specifically 8.00, 8.25, and 8.50 days post-fertilization, to focus on cells with the global cell-type annotation “gut”. These cells correspond to maturing gut cells that were demonstrated to have distinct marker genes that can indicate differentiation between different cell types. Gene expression was quantified using *alevin* run in its default mode, which incorporates multi-mapping reads via the EM algorithm, with 20 bootstrap replicates additionally generated to obtain the mean and variances for compressed uncertainty analysis. We additionally ran *alevin* without the EM step by using the `--noem` flag, which discards multi-mapping reads and thus provides quantification results more comparable to *dropEst* or *Cell Ranger*.

The analysis of cells at 8.00, 8.25, and 8.50 days post-fertilization involved 20,401 cells, and we randomly chose 500 from each time point to include in the trajectory analysis. The subsetting was performed to incorporate cells from each time point that were distributed along the entire developmental trajectory while ensuring computational scalability for the results run on the pseudo-inferential replicates. Trajectory-based differential expression analysis was conducted using the procedure discussed in section 2.4. Hypothesis testing was conducted using the `associationTest` from *tradeSeq* to test for general differences in expression across lineages. We ran the procedure on the counts from *alevin* that incorporate multi-mapping reads using the EM algorithm, and repeated the analysis on the counts that do not incorporate multi-mapping reads and were generated without using the EM algorithm. We additionally simulated 20 pseudo-inferential replicates from the negative binomial distribution using the procedure described in Section 2.3, and combined results across replicates using the procedures described in Section 2.4. Clustering assignment of cells and estimated pseudotimes and lineages were fixed to be those estimated from the EM count point estimates in all cases to ensure all results could be compared as directly as possible.

## 3 Results

### 3.1 Disk space and memory comparison

We first compared the total disk space (in GB) required to store the full object output by *tximport* for the trajectory simulations in a gzip compressed binary format as well as the total memory required (in GB) to load the object in *R* with and without including 20 bootstrap replicates across 100 and 250 cells in Table S1. Matrices within the object are stored in a sparse format, greatly reducing disk space and memory required to load the object into *R*. However, both disk space and memory required to load the object into *R* increased approximately linearly with the number of cells, and storage and memory requirements for results without bootstrap replicates are approximately 18% and 14% of the amounts required for results including all 20 bootstrap replicates. Especially given recent advances in scRNA-seq technology that have made it possible that a single experiment could comprise many thousands or even millions of cells (Lähnemann *et al*., 2020), the disk space and memory required to store results that include all bootstrap replicates may become intractable.

### 3.2 Coverage

Coverage of each interval type was evaluated using the data from the two group difference simulation, stratifying by InfRV and expression level. The InfRV measure is discussed in Section 2.3, but briefly it is a numeric measure that quantifies inferential uncertainty that is roughly stabilized across the range of the counts (Zhu *et al*., 2019). Results show nearly identical coverage values between the two interval types, indicating storage of the sample mean and sample variance of the bootstrap replicates is sufficient to capture the gene-level inferential uncertainty present across the replicates (Figure 2). Coverage tended to be lower for some genes in the upper 10% of InfRV level that are not in the upper 10% of expression level. Interval width tended to be larger for genes in the upper 10% of expression and for genes in the top 10% of InfRV (Figure S1). The distributions of interval widths were nearly identical between the two interval construction methods.

**Figure 2:**
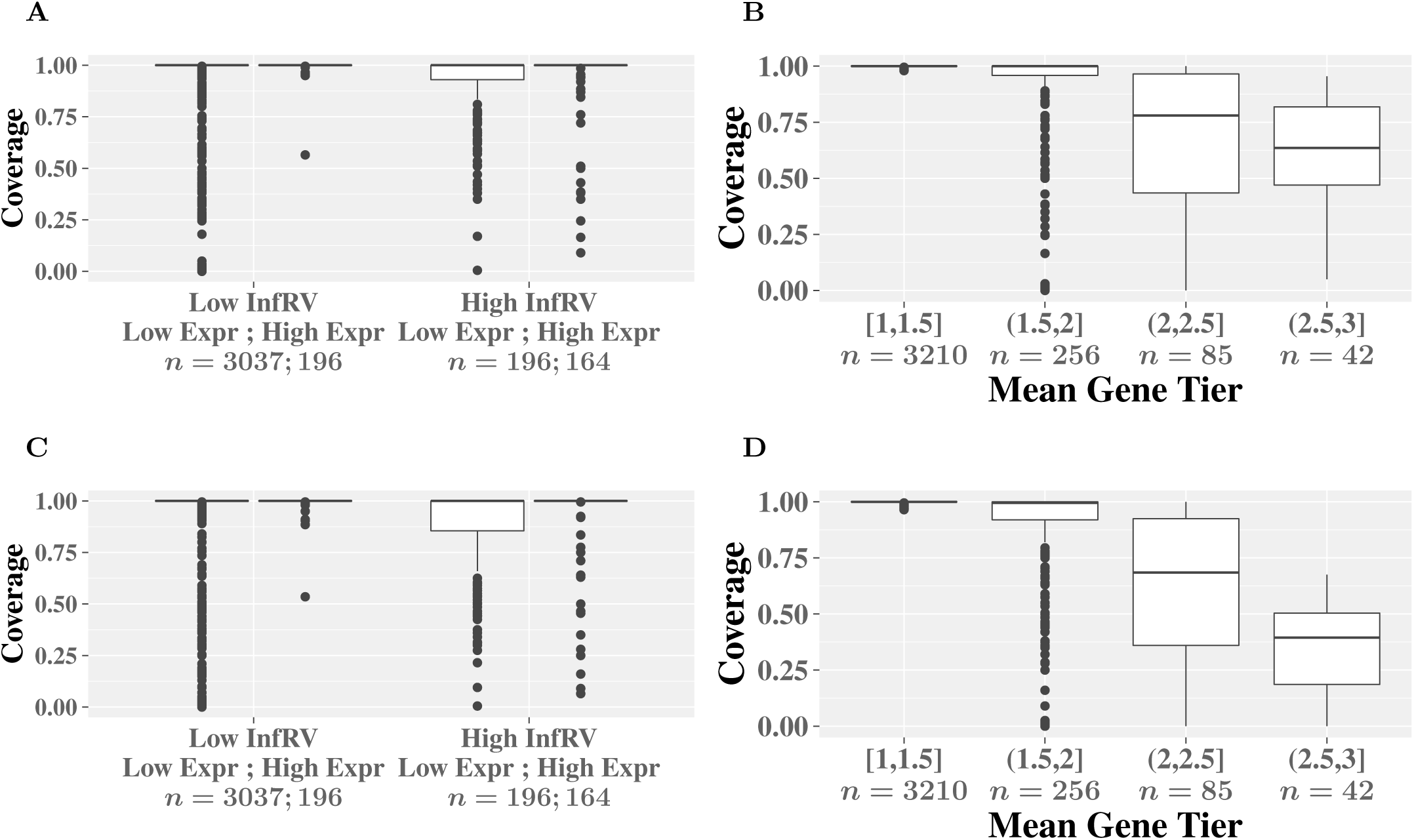
Per-gene coverage comparisons for the 95% intervals calculated using negative binomial distribution quantiles (A and B) and quantiles from the bootstrap empirical distribution (C and D), for the two group difference simulation. Panels A and C are stratified by inferential uncertainty (InfRV) and expression level, while panels B and D are stratified by the average gene tier value across samples. “High” InfRV and expression correspond to the top 10% of InfRV and gene-level counts and respectively.

Coverage of each interval type was also evaluated across the level of uniqueness in the reads contributing to the gene’s expression, as recorded in the gene-by-cell “tier” information output by *alevin* (Srivastava *et al*., 2019). A specific gene and cell combination is assigned a tier value ranging from 0 to 3, with a value of 0 indicating no reads from a cell mapped to the gene, 1 indicating that the gene had *some* unique reads (either all, or a mix of unique and ambiguous), 2 indicating the gene had only ambiguous reads but appeared in a multi-mapping network in which other genes had uniquely mapping reads, and 3 indicating the gene itself, and all other genes in its multi-mapping network, had only ambiguous reads. The overall tier value for a gene was computed as the average of all cell-specific tier values that are greater than zero to ensure cells with no reads mapping to a particular gene did not affect the gene’s overall tier rating. Coverage decreased as the overall tier value increased, corresponding to lower overall uniqueness in the reads contributing to the gene’s expression across cells (Figure 2). Coverage was nearly identical across the two interval types, again indicating storage of the mean and variance was sufficient to capture the gene-level inferential uncertainty present in the bootstrap replicates. The median widths of the intervals decreased as gene tier increased past 2 (fewer unique reads for quantification) but did not differ appreciably between the two interval construction methods (Figure S2). Similar plots using a gene’s uniqueness ratio, which is the proportion of *k*-mers of length 31 present in any of the gene’s transcripts that are not shared with any other genes (Srivastava *et al*., 2019), are given in Figures S3 and S4. Coverage decreased as the sequences contributing to the gene became less unique but the width of the intervals did not change appreciably across gene uniqueness.

We additionally evaluated gene-specific coverage performance across cells in Figures S5, S6, S7, and S8. Coverage of the simulated count varied greatly across genes and cells, with Figure S5 demonstrating low coverage across cells, Figures S6 and S7 demonstrating very high coverage across cells, and Figure S8 demonstrating more variation in coverage across cells. However, coverage results were again very similar between the two interval types.

To evaluate the impact of gene filtering on coverage, we replicated the gene tier coverage plots using all 57,111 genes that were able to be used across the simulation pipeline (Figure S9). Coverage tended to be higher than the corresponding results that filtered genes (Figure 2), indicating lowly expressed counts tended to be easier to cover with intervals than more highly expressed ones. This was further confirmed by removing all counts of 0 from the coverage evaluation for all 57,111 genes, which resulted in significantly lower coverage for genes with high overall tier values (Figure S10). Additionally, coverage results presented in Figure 2 did not differ appreciably when using 100 bootstrap replicates instead of 20 (Figure S11).

Lastly, we evaluated coverage using the simulated trajectory counts from the *dynverse* framework. Results from the trifurcating trajectory simulation with 100 cells are presented in Figures S12 and S13. Coverage from this simulation tended to be significantly lower than the coverage for the two group difference simulation presented in Figure 2 for genes in the upper 10% of quantification uncertainty. This was likely because the expression levels across genes are significantly higher than is typically present in real datasets, with nearly 50,000 genes being highly expressed enough to pass filtering for this simulation.

### 3.3 Trajectory-based differential expression analysis

We used *tradeSeq* to evaluate the effect of incorporating quantification uncertainty into trajectory-based differential expression analysis using pseudo-inferential replicates. Using only the *alevin* point estimates of abundance generally resulted in high sensitivity and often conserved the desired FDR threshold. However, incorporation of quantification uncertainty resulted in reduced FDR, particularly for genes in the upper 20% of InfRV. This was especially true for the `startVsEndTest` and `patternTest` results for the 100 cell trifurcating trajectory simulation. Results for these two tests are shown in Figures S14 and S15 respectively, while results for the `associationTest` and `diffEndTest` are shown in Figures S16 and S17 respectively. Sensitivity when using the mean statistic and Pval50Perc approaches was comparable to use of the point estimates. An example of how the incorporation of quantification uncertainty can benefit analysis can be seen in the `startVsEndTest` results, where use of the point estimates of counts for genes within the highest InfRV category resulted in 8% observed FDR at a nominal 5% FDR, while the three uncertainty-incorporating methods all had observed FDR less than nominal 5%. Additionally, results analogous to Figures S14 and S15 run on the actual bootstrap replicates from *alevin* are shown in Figures S18 and S19. These significance results are nearly identical to results discussed above, indicating that use of pseudo-inferential replicates generated from a negative binomial distribution in place of the actual bootstrap replicates results does not significantly impact downstream results. Results analogous to Figures S14 and S15 run using 100 simulated pseudo-inferential replicates did not differ substantially from results with only 20 (Figures S20 and S21). This indicated 20 pseudo-inferential replicates were sufficient to incorporate quantification uncertainty into the analysis. Lastly, our proposed Pval50Perc and Pval75Perc approaches showed very similar performance to the similar procedure motivated from *RATs* (Froussios *et al*., 2019) that conducts FDR correction *before* selecting the 50th or 75th percentile of adjusted *p*-values as the final value instead of performing the FDR correction *after* selecting the final raw *p*-values (Figures S22 and S23).

To illustrate the advantages of quantification uncertainty on particular genes, we focused on 15 null genes that had a mean count *>* 5 across cells and had high inferential uncertainty (average InfRV *>* 0.5). *P* -values for these genes from the `startVsEndTest` for results calculated using the *alevin* point estimates as well as for Pval50Perc and Pval75Perc are respectively plotted in Figures S24 and S25. Use of the inferential replicates eliminated false positives at the 0.01 FDR level: use of Pval50Perc eliminated 7 of 15 false positives, while use of Pval75Perc eliminated 10 of 15 false positives. Pval75Perc correctly shifted the *p*-value towards 1 for all cases, while Pval50Perc shifted the *p*-value towards 1 in every case except one.

The 250 cell trifurcating trajectory simulation also showed reduced FDR levels but the FDR from the *alevin* point estimates was lower in this simulation than for the 100 cell simulation, meaning less improvement in the FDR from incorporation of quantification uncertainty was possible. We interpret this to be indicative of increased accuracy in the pseudotime and lineage estimation relative to the 100 cell case, resulting in quantification uncertainty having less impact on final significance results across all genes. Results for the 250 cell trifurcating lineage simulation are given in Figures S26, S27, S28, and S29. Significance results for the bifurcating lineage simulation showed similar patterns to results from the trifurcating trajectory simulation, with the FDR always being reduced by incorporating quantification uncertainty via inferential replicates (data not shown). However, the improvements were smaller than those present in the trifurcating trajectory, indicating quantification uncertainty had a smaller effect on the final significance results than for the trifurcating trajectory case. Two dimensional principal component plots of each cell across known pseudotimes for the 100 cell and 250 cell trifurcating simulations are given in Figures S30 and S31 respectively, with the fit lineages from *slingshot* being plotted using the black lines.

### 3.4 Swish with pseudo-inferential replicates

We additionally evaluated and compared *Swish* with the proposed *splitSwish* function. Load time, compute time, and memory comparisons are given in Table 1 and demonstrate that usage of *splitSwish* instead of *Swish* was able to greatly reduce the size of the quantification object and memory required to complete the analysis. Compute time summing across all eight jobs was increased with *splitSwish* compared to *Swish*, but per job the compute time was reduced about four-fold. Sensitivity and false discovery rates were comparable between *splitSwish* and *Swish* (Figure S32).

**Table 1:** Computation comparisons for *Swish* and *splitSwish* for the two group difference simulation. Results include 60,179 genes across 200 cells, with 20 bootstrap replicates for *Swish* and 20 pseudo-inferential replicates for *splitSwish*. R object size and load time differ across methods, as *Swish* uses full bootstrap replicate matrices while *splitSwish* uses compressed inferential uncertainty. Max memory and compute time are provided per job (*n* = 8) for *splitSwish*.

| Method | R object size (MB) | Max memory (GB) | Load (s) | Compute (s) |
| --- | --- | --- | --- | --- |
| *Swish* | 853 | 4.90 | 28.2 | 78 |
| *splitSwish* | 138 | 1.08 | 1.5 | 20 |

### 3.5 Mouse embryo data

We lastly evaluated the impact of multi-mapping reads and quantification uncertainty on a trajectory-based differential expression analysis of mouse embryo data collected at 8.00, 8.25, and 8.50 days post-fertilization. We found that counts incorporating multi-mapping reads can differ greatly from those that do not for certain genes while being virtually unchanged for other genes. This was even true for genes within a common gene family, where counts for certain genes within the family were significantly underestimated without incorporating multi-mapping reads.

For example, the Nme1 and Nme2 genes are known to be part of the Nm23 gene family, and have been shown to be responsible for the majority of NDP kinase activity in mammals (Postel *et al*., 2009) along with other cellular processes (Boissan *et al*., 2018). Nme1 and Nme2 can be co-transcribed, forming a fusion protein (Akiva *et al*., 2006; Prakash *et al*., 2010). Mice that had both genes deleted have been previously found to suffer stunted growth and die perinatally (Postel *et al*., 2009), demonstrating the clear importance of the gene family in mammalian development. The gene family has additionally been shown to play a vital role in non-mammal vertebrate species (Desvignes *et al*., 2009), and low expression of Nm23 has long been identified to play crucial role in cancer mestasis in humans (MacDonald *et al*., 1995; Hartsough and Steeg, 2000; Jarrett *et al*., 2013).

In the mouse embryo dataset, a comparison of Nme1 and Nme2 counts estimated with and without the EM algorithm (henceforth referred to as “EM” and “no EM” respectively) are presented in Figure 3. Counts for Nme1 were nearly identical whether incorporating multi-mapping reads or not, resulting in the predicted counts across pseudotime for each lineage having similar shapes with and without incorporating multi-mapping reads (Figure S33). In contrast, counts for Nme2 were found to be much lower and near zero without incorporating multi-mapping reads, resulting in the predicted counts across pseudotime for each lineage being much lower when ignoring multi-mapping reads (Figure S33).

**Figure 3:**
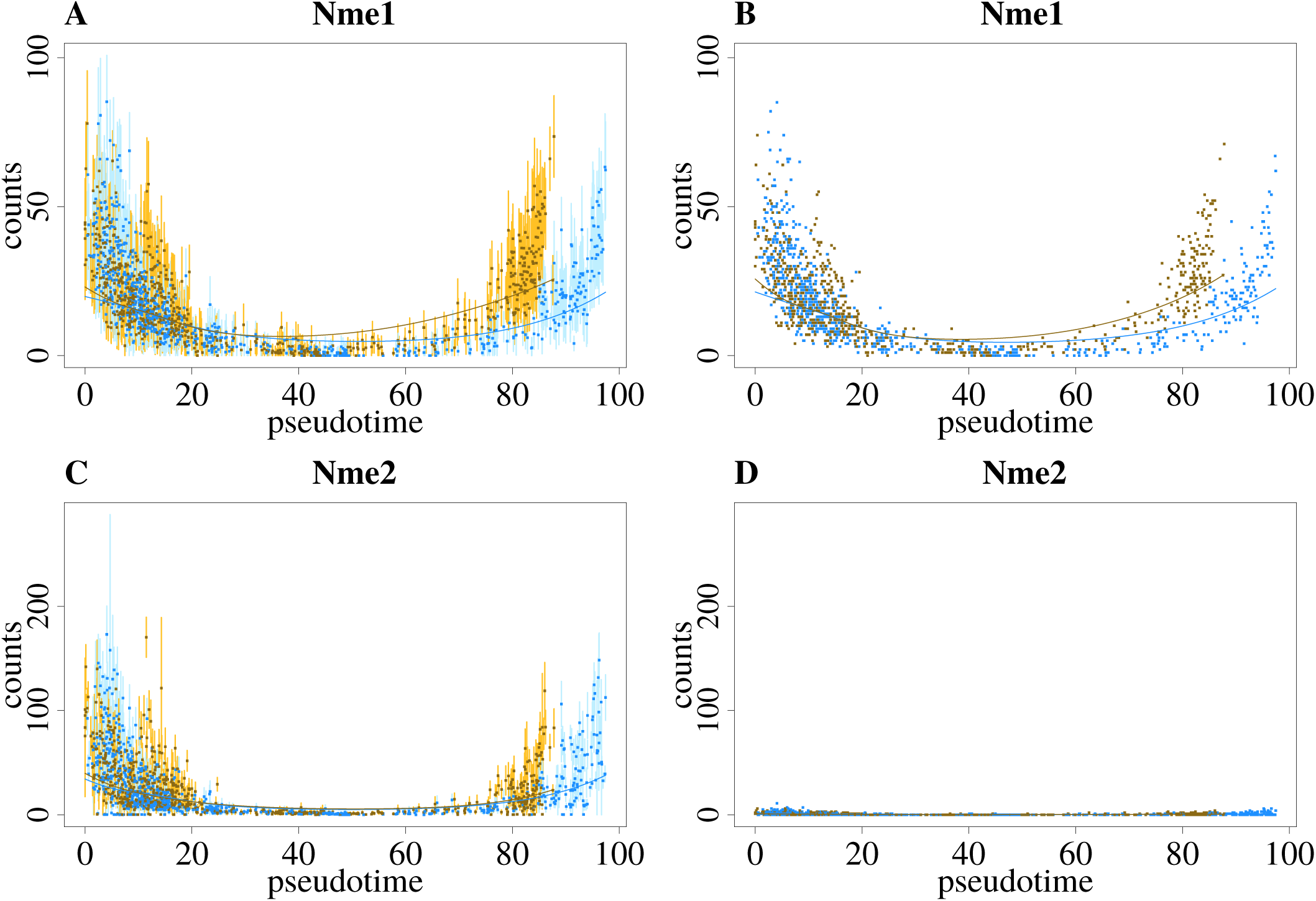
Comparison of counts across pseudotime for Nme1 and Nme2 for counts generated incorporating multi-mapping reads using the EM algorithm (A and C) and without incorporating multi-mapping reads (B and D). Counts are colored according to assignment to one of two lineages. Points represent mean of bootstrap replicates and vertical bars represent 95% normal-based intervals in A and C, while points in B and D provide estimated counts. Curves plot the fitted GAMs across pseudotime for each lineage.

Pseudo-inferential replicates and the proposed Pval50Perc method were used to conduct significance testing for the uncertainty-aware trajectory analysis (“EM with uncertainty”). Adjusted *p*-values from the `associationTest` were highly significant (*<* 10^*−*12^) for all three scenarios (“EM”, “EM with uncertainty”, “no EM”) for Nme1 and Nme2 but a manual inspection of the fit GAMs in Figure S33 for the “no EM” results revealed the predicted counts nevertheless do not differ to a large extent across pseudotime. This would lead to the incorrect conclusion that Nme2 was always very lowly expressed across pseudotime, despite a statistically significant association with pseudotime. Very similar results were found when the fit lineages and resulting GAMs for the “no EM” results were allowed to differ from those fit for the “EM” results (Figure S34).

Additional examples of genes with much lower estimated counts when ignoring multi-mapping reads include Hmgb1 and Rpl36a (Figures S35 and S36). There were large differences in the fit GAMs for both genes for the “EM” and “no EM” results (Figure S37), though *p*-values from the `associationTest` were all highly significant (*<* 10^*−*12^) for both genes for all three scenarios. Similar differences in estimated counts when not incorporating multi-mapping reads were also present for 358 genes when subsetting based on the total gene count across cells being more than 50% higher or lower across quantification method (Figure S38).

## 4 Discussion

Previous work had demonstrated the necessity of incorporating multi-mapping reads into scRNA-seq analysis, as discarding them could result in up to a 23% decrease in the number of reads used for quantification (Srivastava *et al*., 2019) and induce systematic bias for certain groups of genes based on coverage and sequence homology. *alevin* incorporates these multi-mapping reads and additionally allows drawing bootstrap replicates to estimate quantification uncertainty that is present due to these multi-mapping reads. Here we demonstrate that storage of the sample mean and sample variance estimates of these bootstrap replicates from *alevin* is sufficient to capture the gene-level inferential uncertainty present in sampled replicates. Pseudo-inferential replicates can be generated from a negative binomial distribution as needed, enabling easier incorporation of quantification uncertainty into downstream analyses. While coverage of the true count does not generally differ with and without compression of quantification uncertainty, certain genes showed very low coverage. Some of these genes showed high levels of quantification uncertainty, but ideally even high quantification uncertainty should not directly result in decreased coverage but instead only larger interval widths. We plan to extend *alevin* to produce posterior Gibbs samples for the underlying Bayesian model. Since Gibbs sampling explores the entire parametric space by fixing other estimates but one, we believe the resulting distribution will represent the uncertainty more accurately than bootstrap sampling. Use of Gibbs sampling would additionally allow constructed coverage intervals to be interpreted as Bayesian credible intervals since a valid posterior distribution would be used in their construction.

A limitation of the compressed uncertainty procedure we have proposed is the fact that it only preserves the marginal gene-level inferential replicate distribution such that it can’t be used with methods that require covariance between pairs of genes or transcripts, such as *terminus* (Sarkar *et al*., 2020). The proposed approach that uses *p*-value quantiles from results repeated across pseudo-inferential replicates to determine significance has the advantage that it can be applied to any statistical method without directly requiring any additional assumptions. The proposed approach that uses the mean test statistic across replicates is similarly flexible but assumes that the mean test statistic follows a parametric null distribution to determine significance. This assumption may not hold in certain situations.

Future work could investigate additional approaches to incorporate quantification uncertainty into downstream statistical analyses and to incorporate uncertainty into additional methods and workflows. Quantification uncertainty has been previously shown to improve performance when incorporated into matrix factorization for microarray analysis (Wang *et al*., 2006) and ordination methods for microbiome analysis (Ren *et al*., 2017; Nguyen and Holmes, 2017), and these and similar methods could be extended to incorporate compressed uncertainty. Future work incorporating uncertainty into trajectory analysis specifically could additionally seek to evaluate the effect of fixing cluster assignments, pseudotimes, and lineages across pseudo-inferential replicates. Keeping these consistent across pseudo-inferential replicates prevents issues that can complicate combination of results across replicates, such as different replicates resulting in a different number of lineages or in different starting and ending clusters. However, this approach will not incorporate uncertainty that manifests itself through differences in cluster assignments, pseudotimes, and lineages themselves.

## Supplementary information

Supplementary results are available in the corresponding Supplement file.

