## Supplementary Tables and Figures for "Compression of quantification uncertainty for scRNA-seq counts"

**Supplementary Table S1:** Comparison of the total file size on disk and memory required in gigabytes (GB) to load the *Alevin* quantification results into *R* using *tximport* with and without 20 bootstrap replicates.

| NumCells | BootReps | SizeOnDisk (GB) | Memory (GB) |
|---|---|---|---|
| 100 | No | 0.08 | 0.25 |
| 250 | No | 0.19 | 0.59 |
| 100 | Yes | 0.43 | 1.77 |
| 250 | Yes | 1.06 | 4.19 |

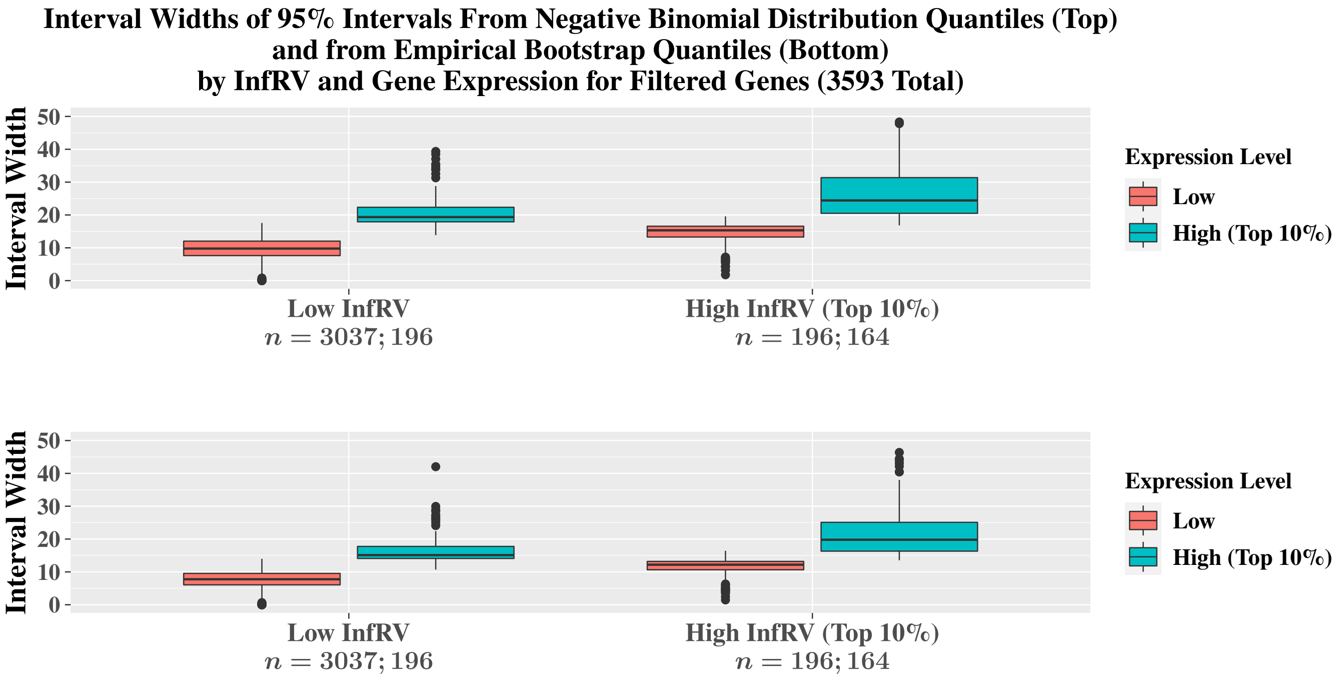


**Supplementary Figure S1:** Coverage comparisons of the 95% intervals calculated using negative binomial distribution quantiles (top) and quantiles from the bootstrap empirical distribution (bottom) stratified by both expression level and InfRV level for the two group difference simulation.

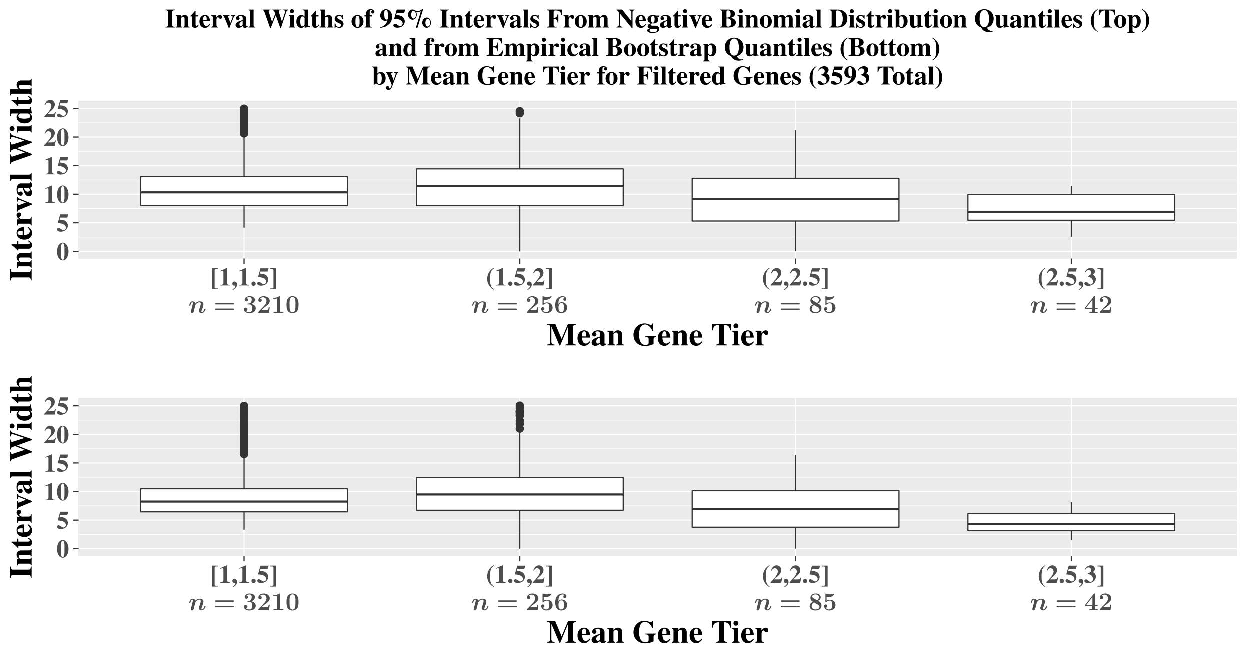


**Supplementary Figure S2:** Coverage comparisons of the 95% intervals calculated using negative binomial distribution quantiles (top) and quantiles from the bootstrap empirical distribution (bottom) stratified by both expression level and InfRV level for the two group difference simulation. Some points are omitted for the [1,1.5] and (1.5,2] gene tier groups.

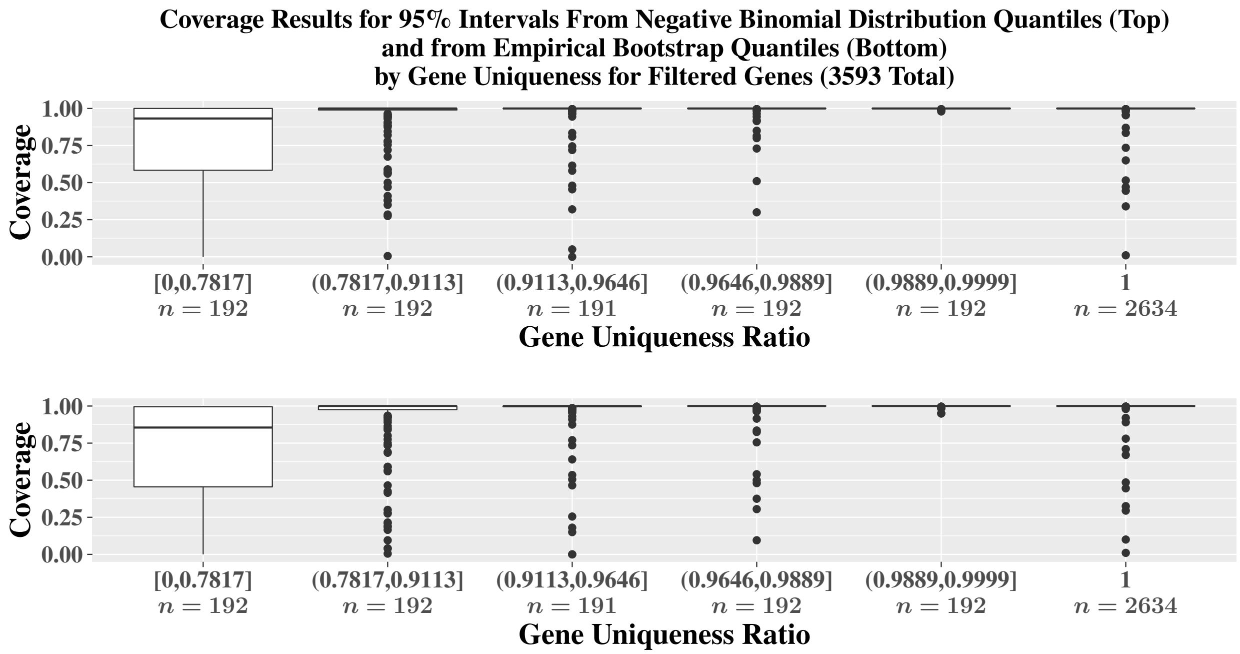


**Supplementary Figure S3:** Coverage comparisons of the 95% intervals calculated using negative binomial distribution quantiles (top) and quantiles from the bootstrap empirical distribution (bottom) stratified by the uniqueness of the gene sequence relative to other genes for the two group difference simulation.

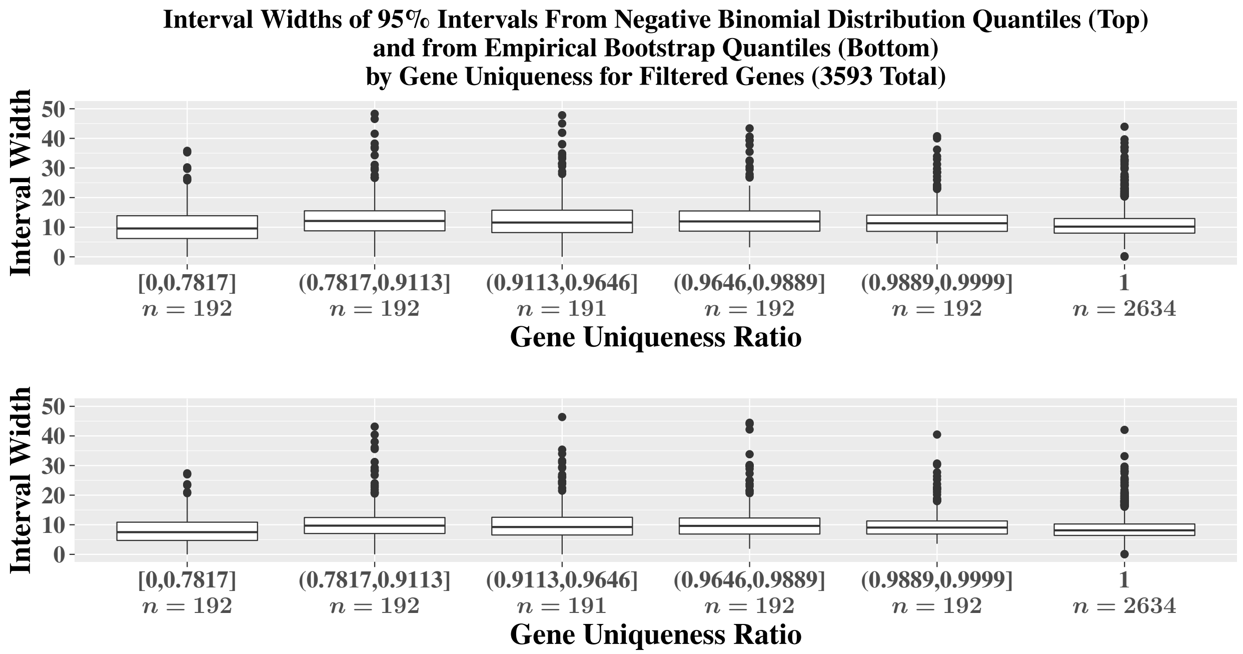


**Supplementary Figure S4:** Comparisons of the widths of the 95% intervals calculated using negative binomial distribution quantiles (top) and quantiles from the bootstrap empirical distribution (bottom) stratified by the uniqueness of the gene sequence relative to other genes for the two group difference simulation.

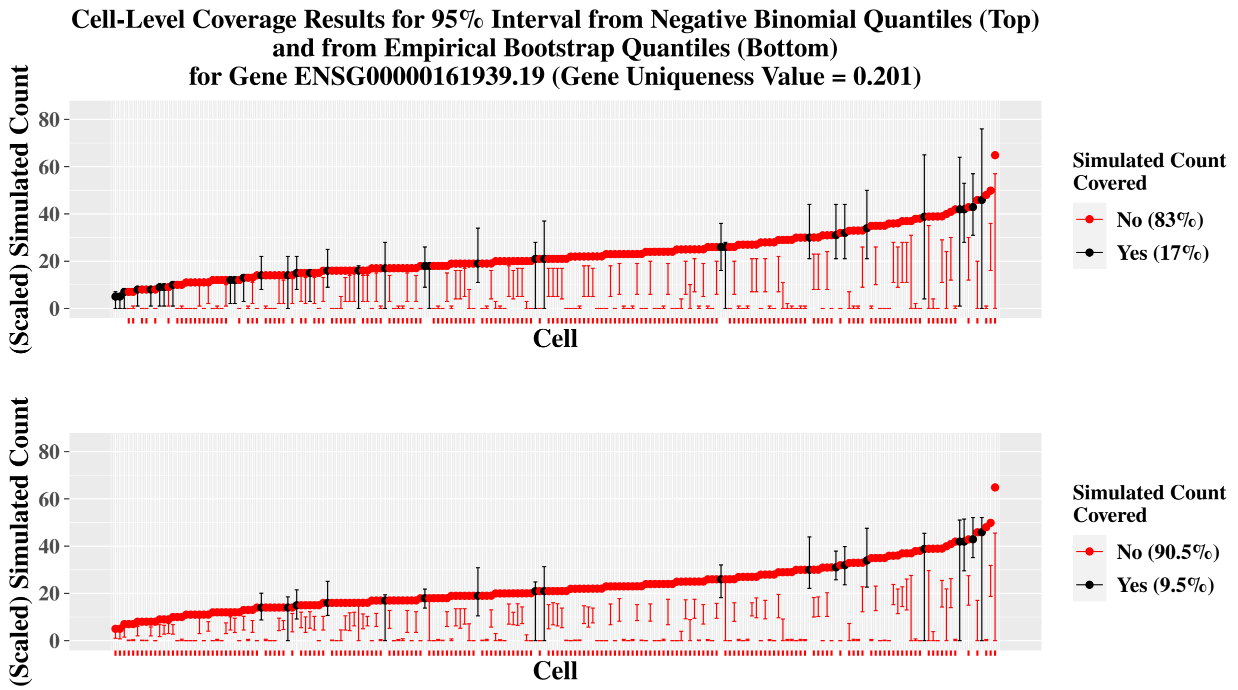


**Supplementary Figure S5:** Comparisons of cell-specific coverages for 95% intervals calculated using negative binomial distribution quantiles (top) and quantiles from the bootstrap empirical distribution (bottom). The points are the simulated count that has been scaled such that the total library size of each cell is equal to the total library size from the *Alevin* quantifications of the simulated reads. The $x$-axis is ordered by the cell-specific simulated count, and the error bars correspond to the upper and lower values of the 95% interval for the cell. Results are from the two group difference simulation.

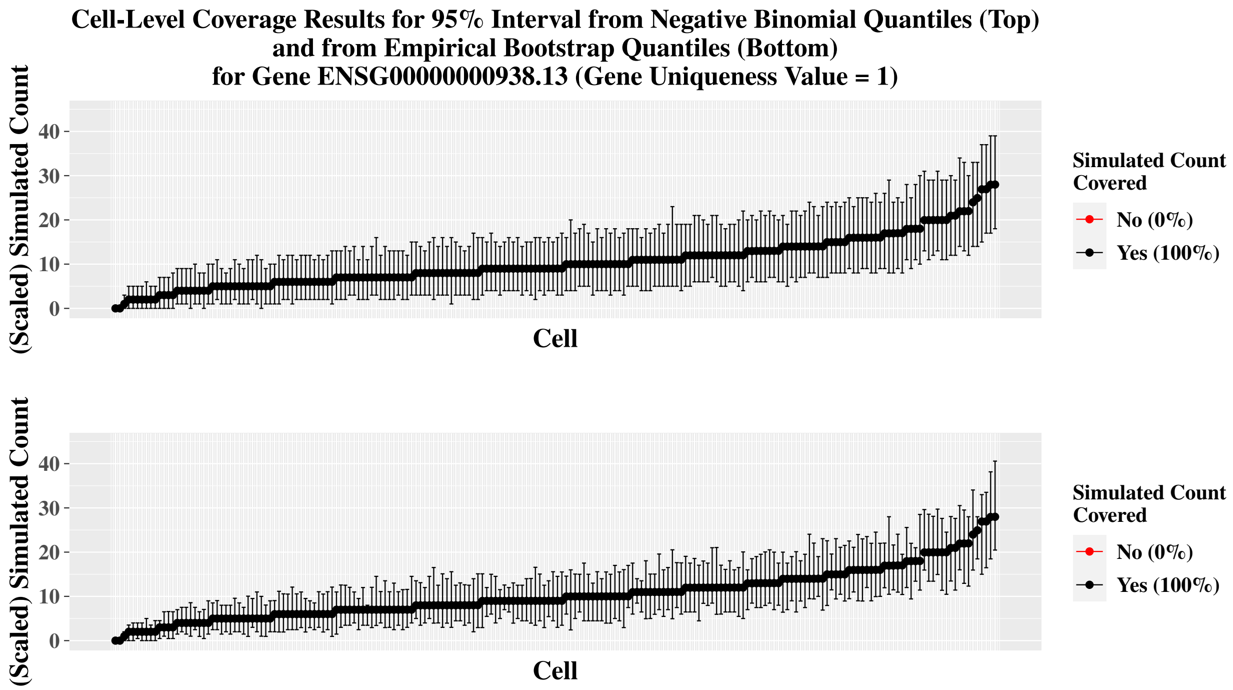


**Supplementary Figure S6:** Comparisons of cell-specific coverages for 95% intervals calculated using negative binomial distribution quantiles (top) and quantiles from the bootstrap empirical distribution (bottom). The points are the simulated count that has been scaled such that the total library size of each cell is equal to the total library size from the *Alevin* quantifications of the simulated reads. The $x$-axis is ordered by the cell-specific simulated count, and the error bars correspond to the upper and lower values of the 95% interval for the cell. Results are from the two group difference simulation.

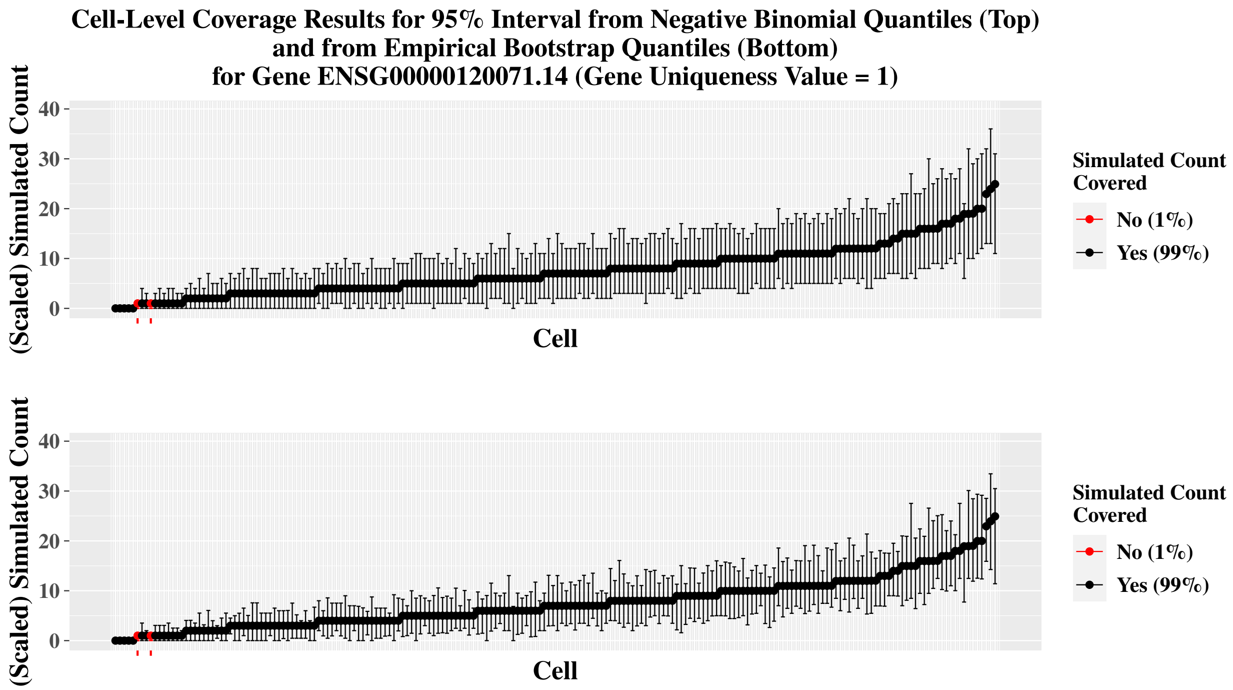


**Supplementary Figure S7:** Comparisons of cell-specific coverages for 95% intervals calculated using negative binomial distribution quantiles (top) and quantiles from the bootstrap empirical distribution (bottom). The points are the simulated count that has been scaled such that the total library size of each cell is equal to the total library size from the *Alevin* quantifications of the simulated reads. The $x$-axis is ordered by the cell-specific simulated count, and the error bars correspond to the upper and lower values of the 95% interval for the cell. Results are from the two group difference simulation.

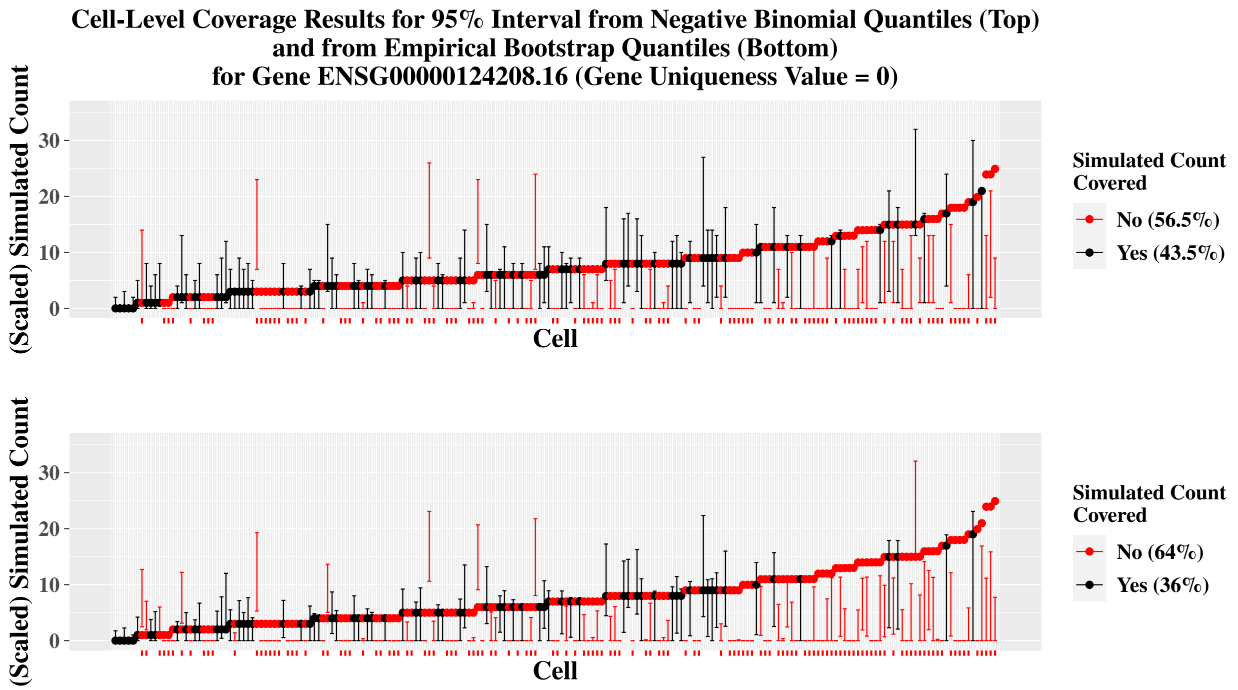


**Supplementary Figure S8:** Comparisons of cell-specific coverages for 95% intervals calculated using negative binomial distribution quantiles (top) and quantiles from the bootstrap empirical distribution (bottom). The points are the simulated count that has been scaled such that the total library size of each cell is equal to the total library size from the *Alevin* quantifications of the simulated reads. The $x$-axis is ordered by the cell-specific simulated count, and the error bars correspond to the upper and lower values of the 95% interval for the cell. Results are from the two group difference simulation.

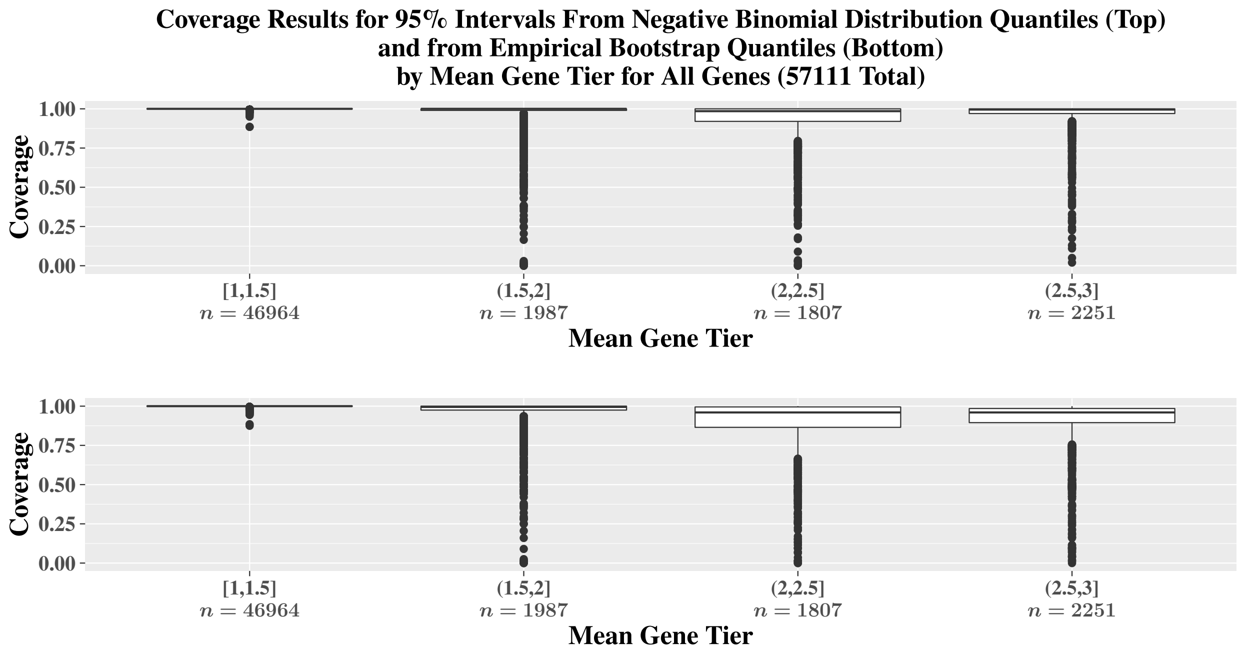


**Supplementary Figure S9:** Coverage comparisons of the 95% intervals calculated using negative binomial distribution quantiles (top) and quantiles from the bootstrap empirical distribution (bottom) stratified by overall tier value of the gene for the two group difference simulation.

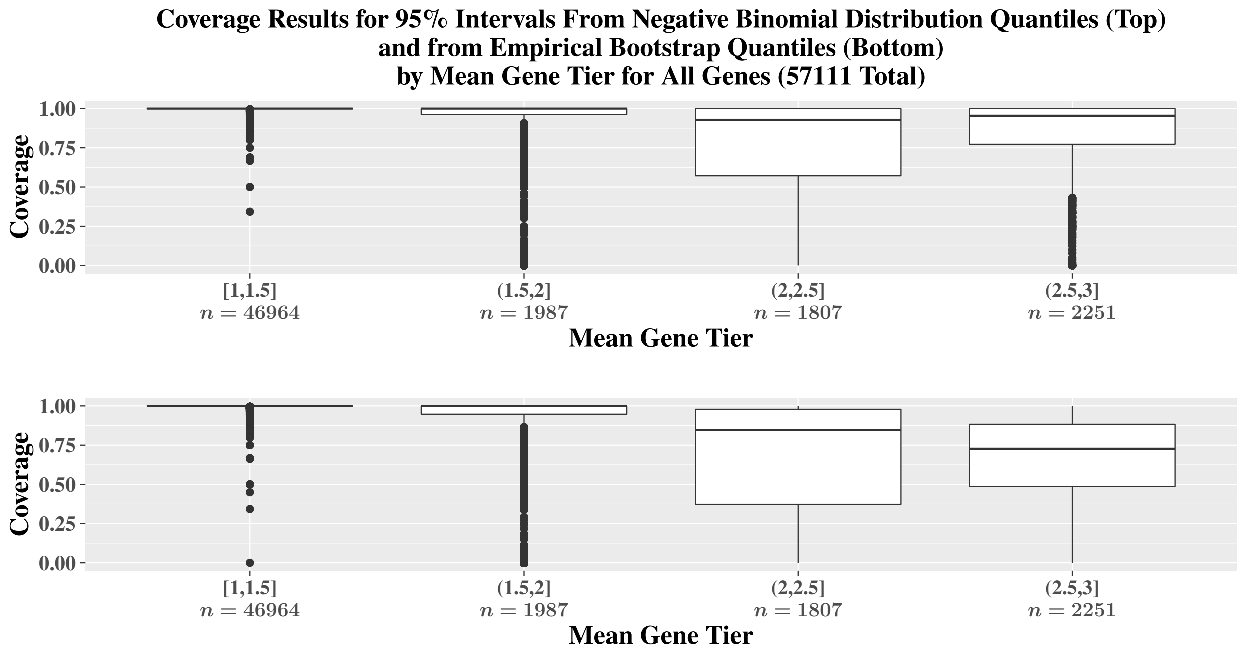


**Supplementary Figure S10:** Coverage comparisons of the 95% intervals calculated using negative binomial distribution quantiles (top) and quantiles from the bootstrap empirical distribution (bottom) stratified by overall tier value of the gene for the two group difference simulation. All zero counts are not included in these results.

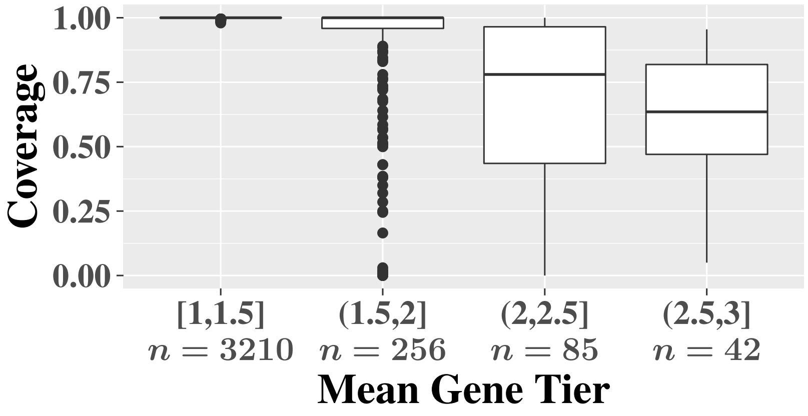


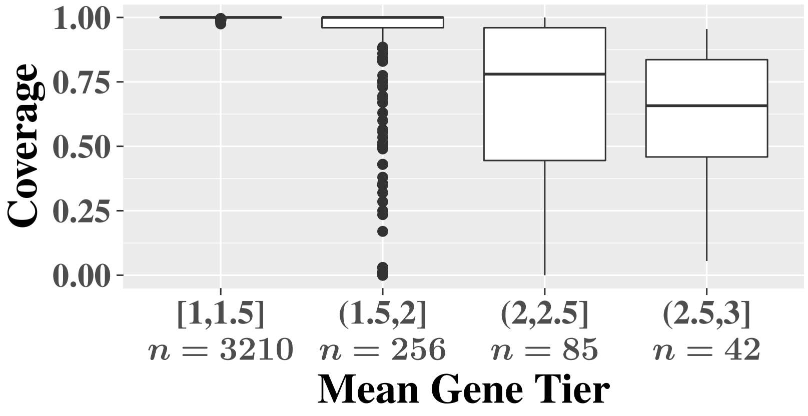


**Supplementary Figure S11:** Coverage comparisons of the 95% intervals calculated using negative binomial distribution quantiles with 20 pseudo-inferential replicates (top) and 100 pseudo-inferential replicates (bottom) stratified by overall tier value of the gene for two group difference simulations.

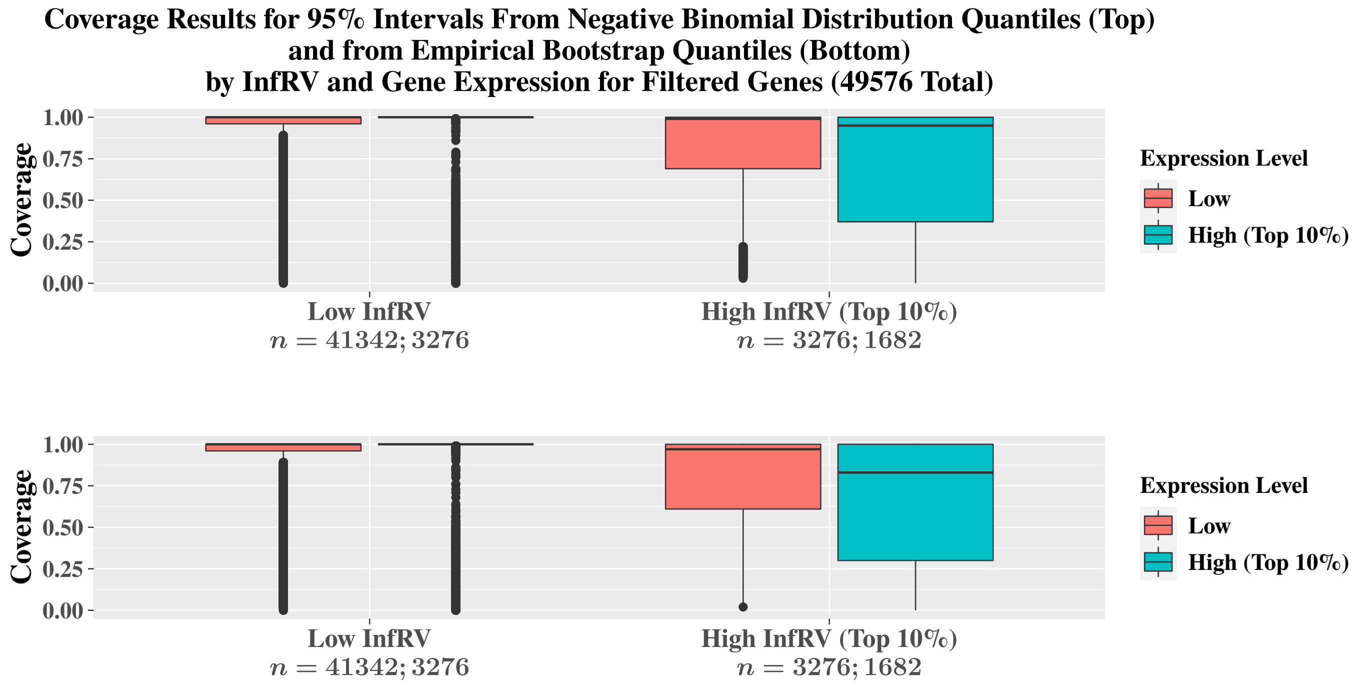


**Supplementary Figure S12:** Coverage comparisons of the 95% intervals calculated using negative binomial distribution quantiles (top) and quantiles from the bootstrap empirical distribution (bottom) stratified by both expression level and InfRV level for the 100 cell trifurcating trajectory simulation.

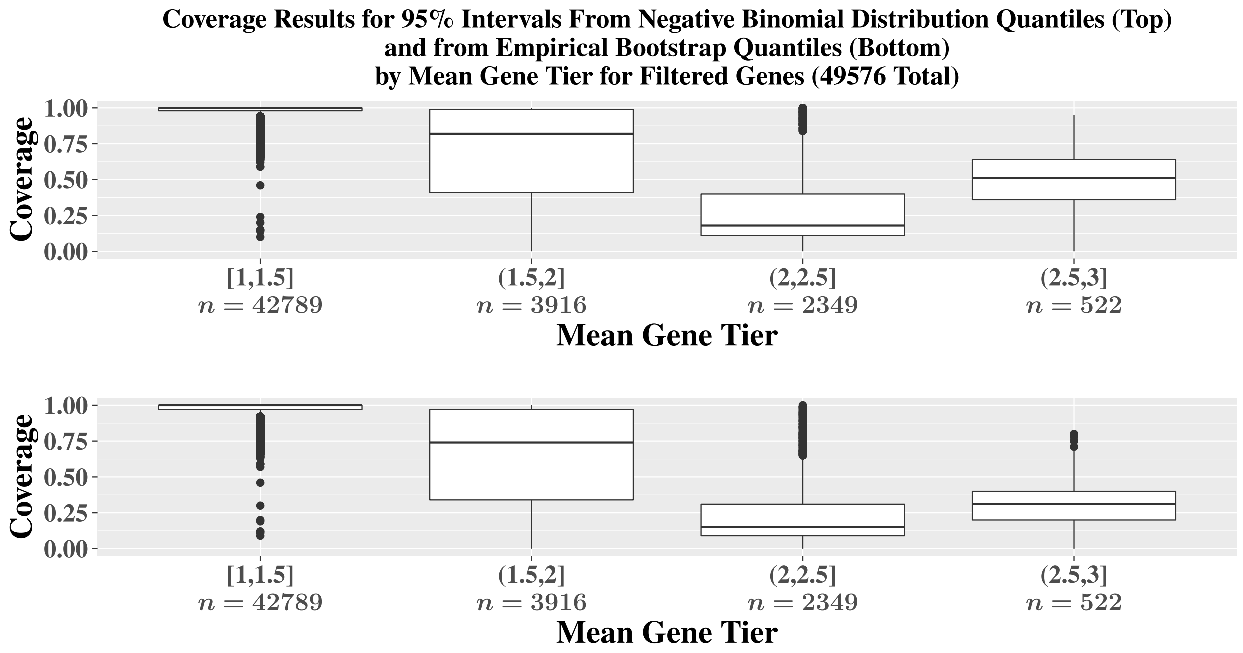


**Supplementary Figure S13:** Coverage comparisons of the 95% intervals calculated using negative binomial distribution quantiles (top) and quantiles from the bootstrap empirical distribution (bottom) stratified by the gene tier value for the 100 cell trifurcating trajectory simulation.

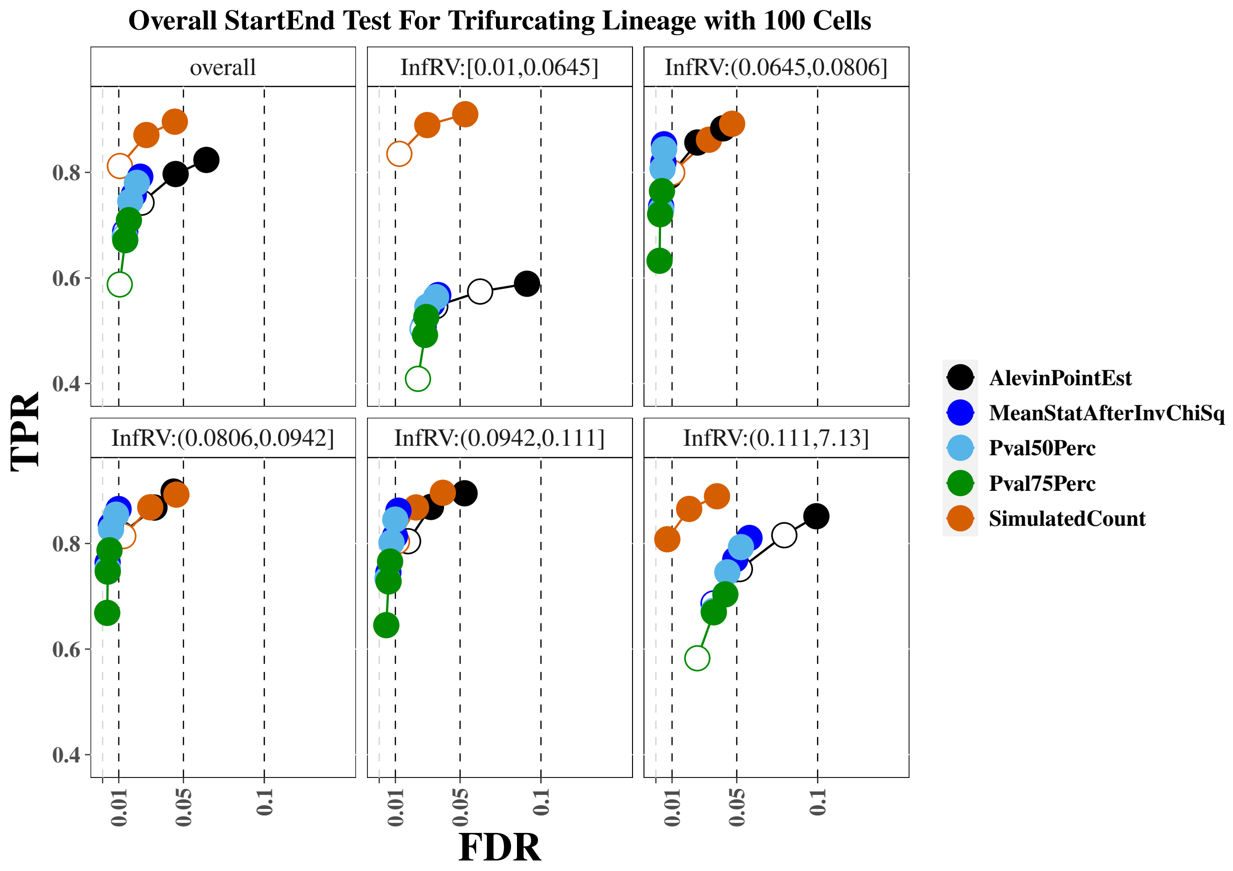


**Supplementary Figure S14:** True positive rate (y-axis) over false discovery rate (x-axis) for the 100 cell trifurcating lineage simulation, using psuedo-inferential replicates. The first panel displays overall performance, while the additional panels stratify genes into fifths based on quantification uncertainty as measured by the *InfRV* statistic averaged across cells. For this and all additional *iCOBRA* plots, the three circles per method indicate performance at nominal FDR cutoffs of 1%, 5% and 10%, with the x-axis providing the observed FDR and the three FDR cutoffs indicated with black vertical dashed lines. A filled circle indicates the desired FDR threshold is conserved, while an open circle indicates the desired FDR threshold is not conserved.

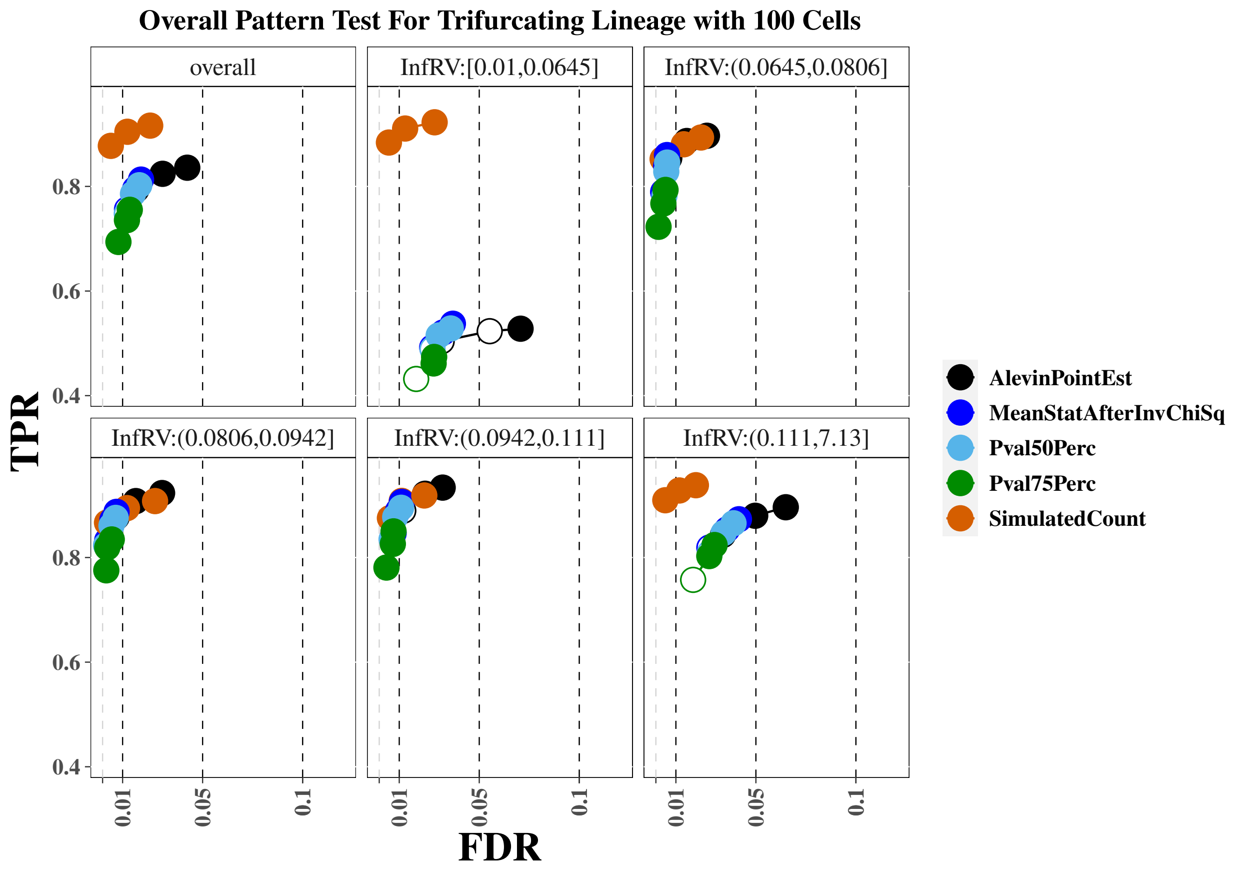


**Supplementary Figure S15:** True positive rate (y-axis) over false discovery rate (x-axis) for the *tradeSeq* for the 100 cell trifurcating lineage simulation, using psuedo-inferential replicates. The first panel displays overall performance, while the additional panels stratify genes into fifths based on quantification uncertainty as measured by the *InfRV* statistic averaged across cells.

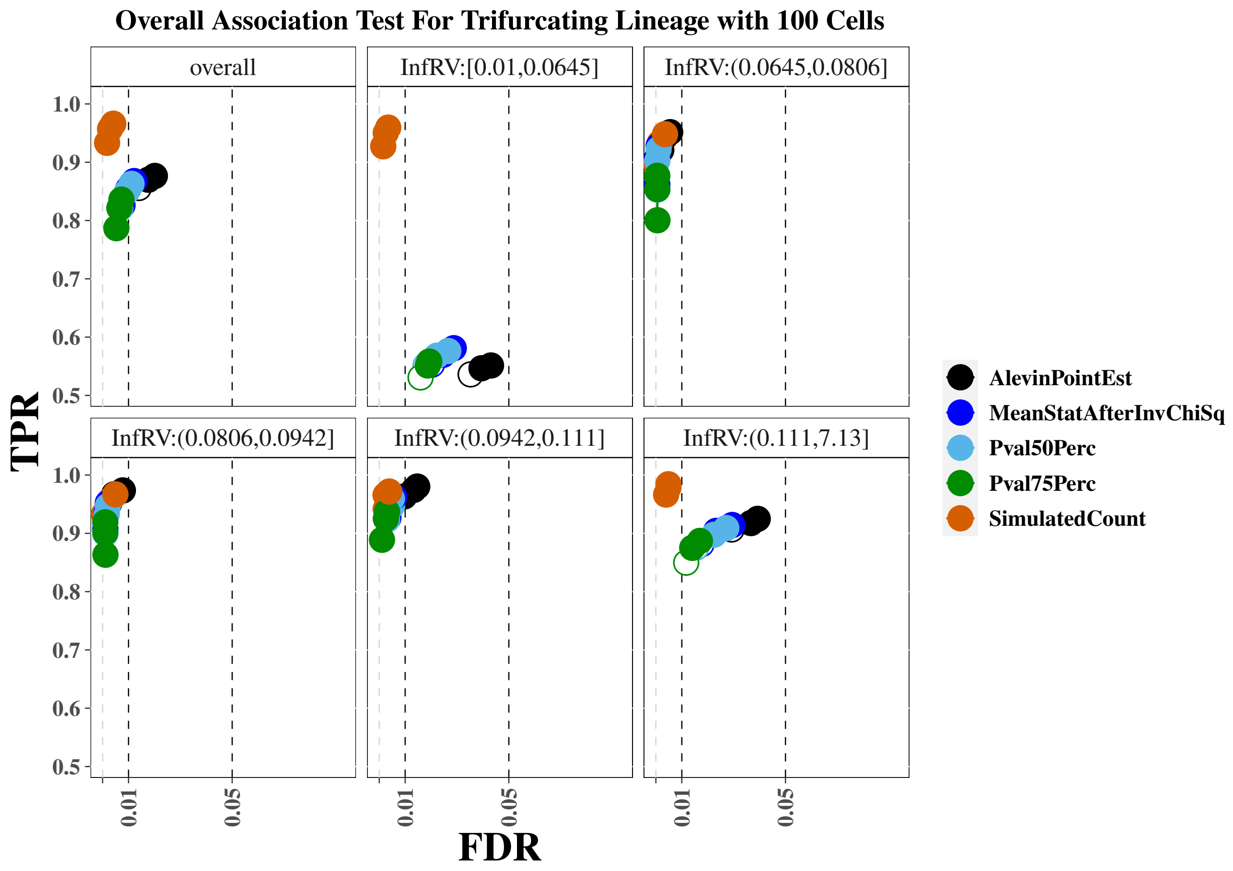


**Supplementary Figure S16:** True positive rate (y-axis) over false discovery rate (x-axis) for the *tradeSeq* for the 100 cell trifurcating lineage simulatio, using psuedo-inferential replicatesn. The first panel displays overall performance, while the additional panels stratify genes into fifths based on quantification uncertainty as measured by the *InfRV* statistic averaged across cells.

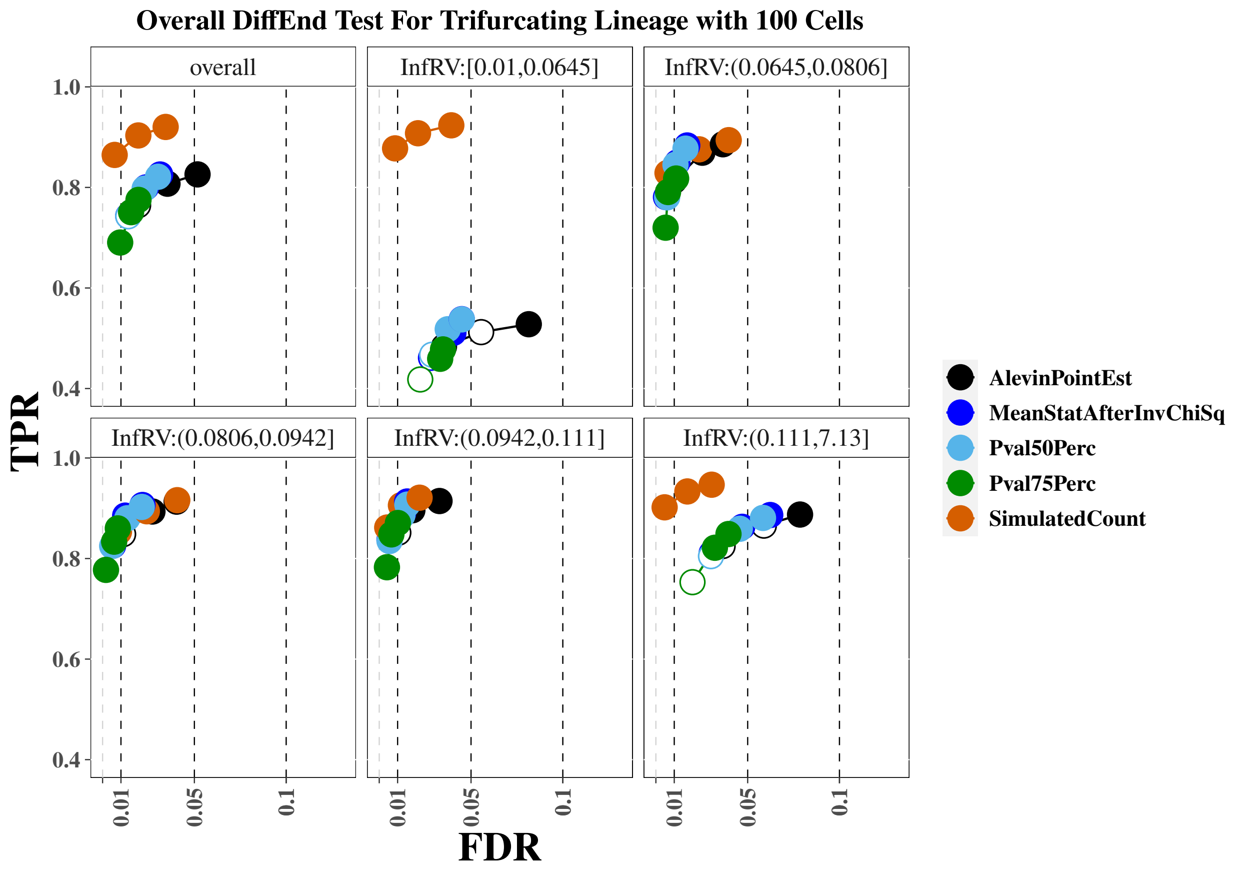


**Supplementary Figure S17:** True positive rate (y-axis) over false discovery rate (x-axis) for the *tradeSeq* for the 100 cell trifurcating lineage simulation, using psuedo-inferential replicates. The first panel displays overall performance, while the additional panels stratify genes into fifths based on quantification uncertainty as measured by the *InfRV* statistic averaged across cells.

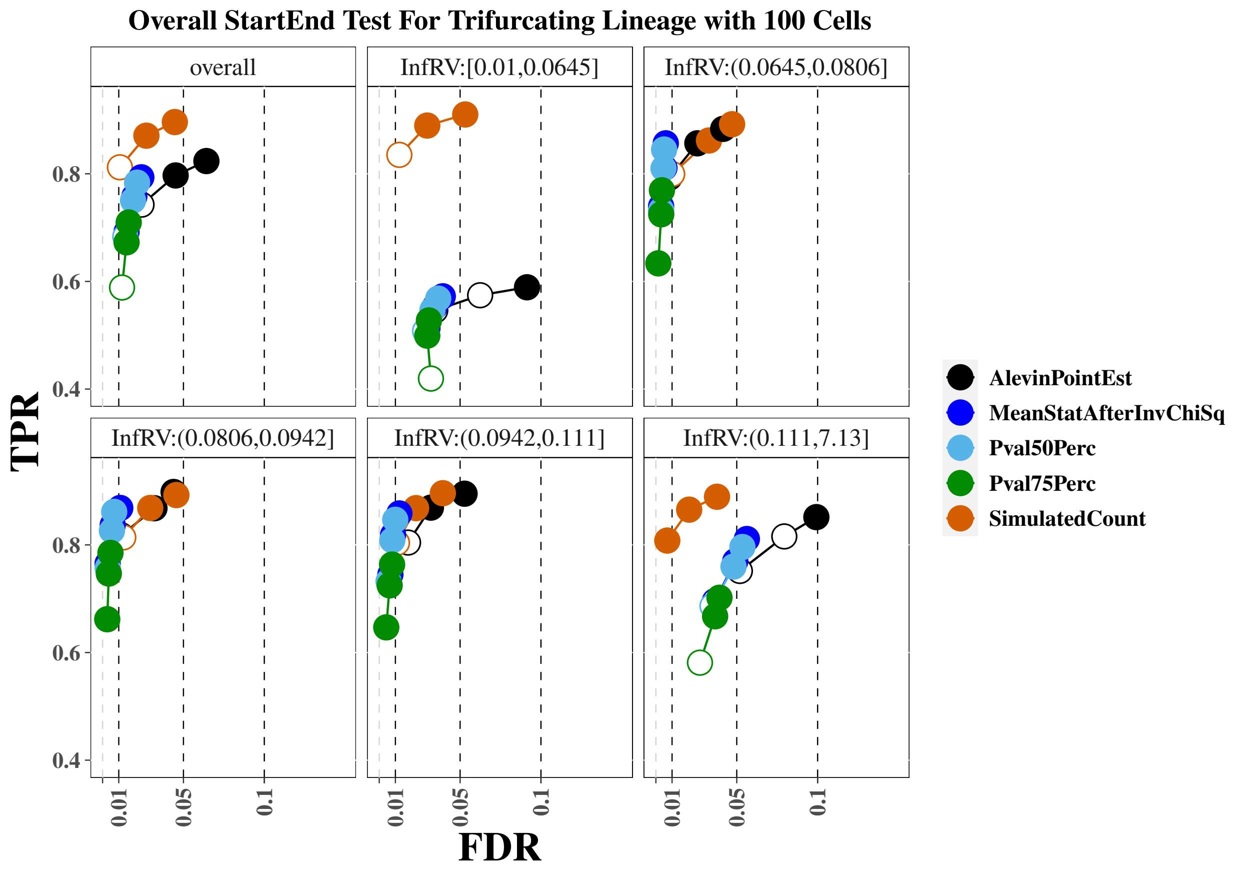


**Supplementary Figure S18:** True positive rate (y-axis) over false discovery rate (x-axis) for the *tradeSeq* for the 100 cell trifurcating lineage simulation. The first panel displays overall performance, while the additional panels stratify genes into fifths based on quantification uncertainty as measured by the *InfRV* statistic averaged across cells. These results were run on actual bootstrap replicates instead of simulated pseudo-inferential replicates, and show very similar performance to results run on the pseudo-inferential replicates.

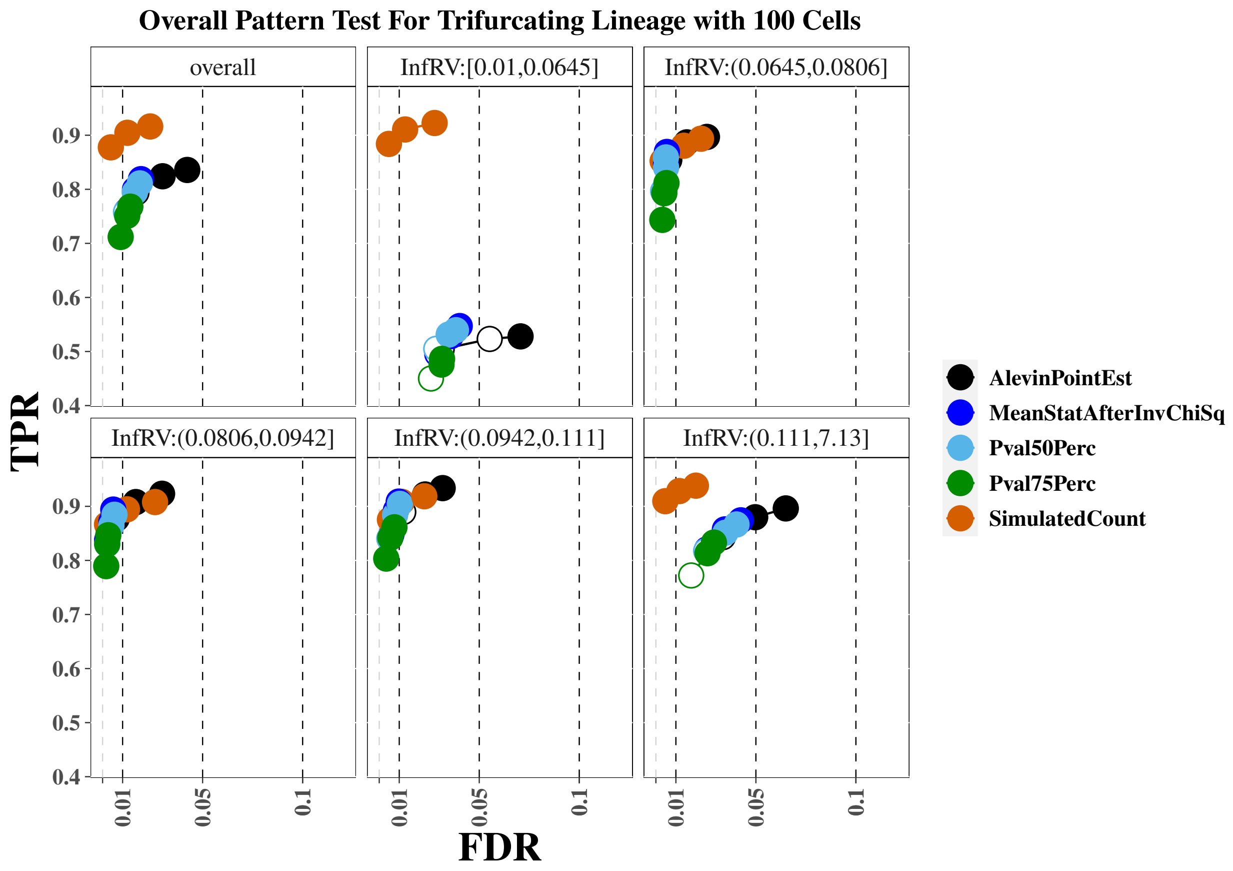


**Supplementary Figure S19:** True positive rate (y-axis) over false discovery rate (x-axis) for the *tradeSeq* for the 100 cell trifurcating lineage simulation. The first panel displays overall performance, while the additional panels stratify genes into fifths based on quantification uncertainty as measured by the *InfRV* statistic averaged across cells. These results were run on actual bootstrap replicates instead of simulated pseudo-inferential replicates, and show very similar performance to results run on the pseudo-inferential replicates.

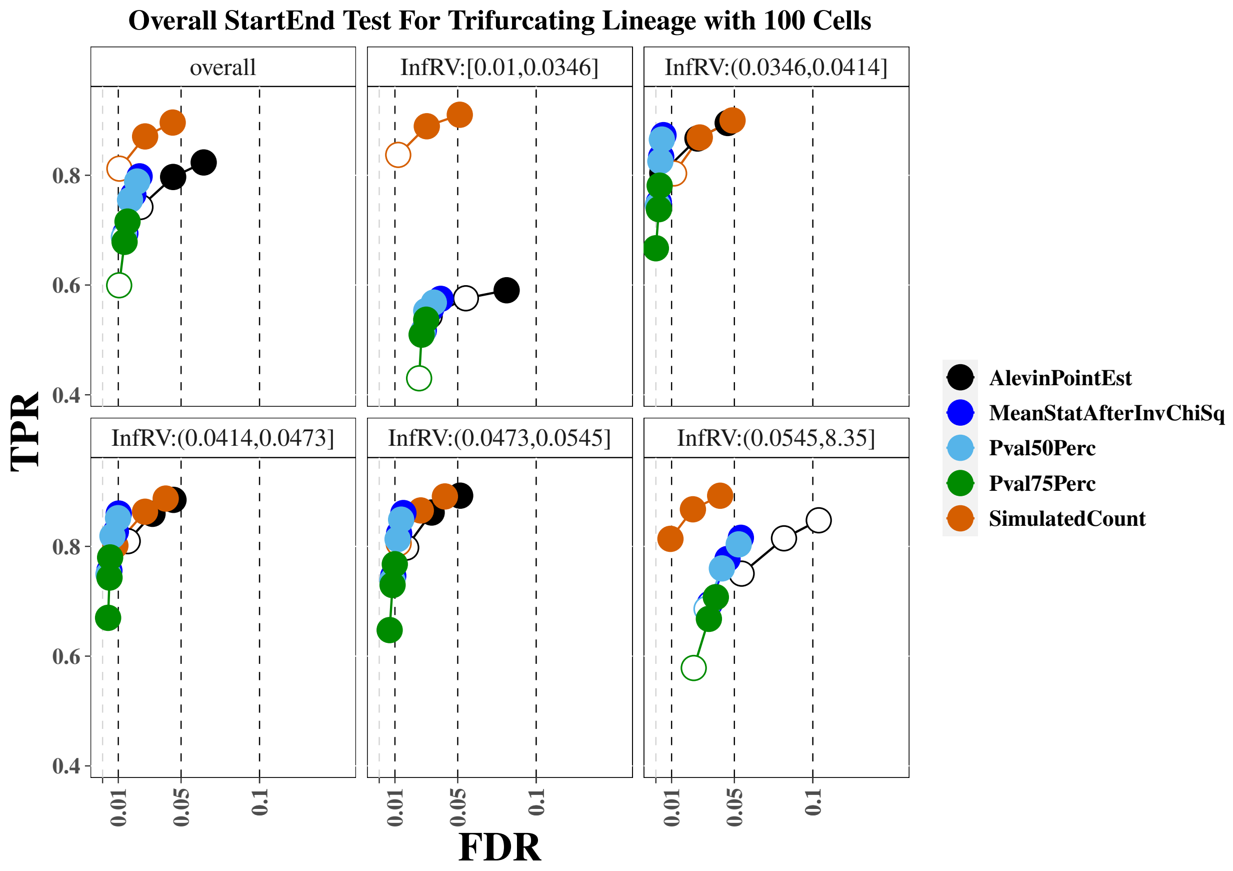


**Supplementary Figure S20:** True positive rate (y-axis) over false discovery rate (x-axis) for the *tradeSeq* for the 100 cell trifurcating lineage simulation. The first panel displays overall performance, while the additional panels stratify genes into fifths based on quantification uncertainty as measured by the *InfRV* statistic averaged across cells. These results use 100 bootstrap replicates instead of 20.

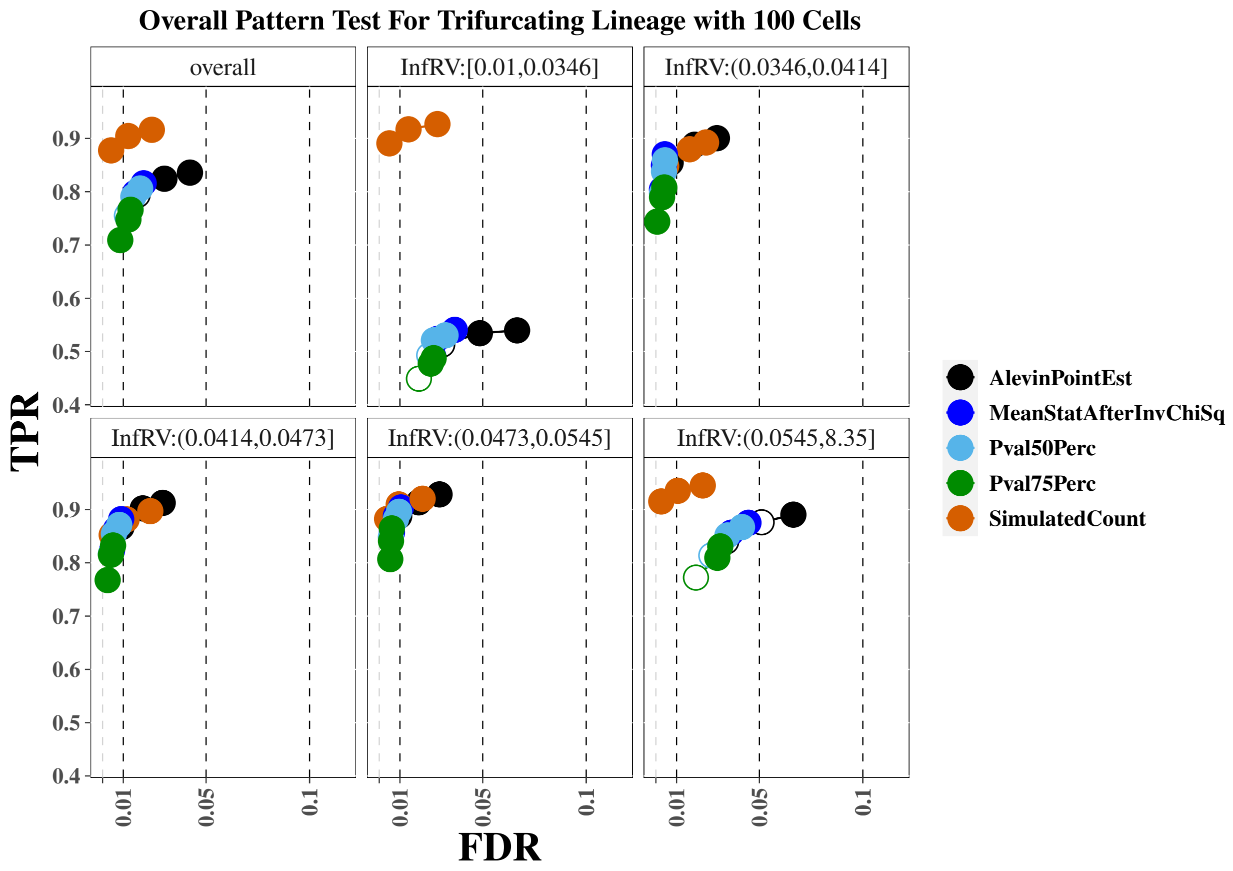


**Supplementary Figure S21:** True positive rate (y-axis) over false discovery rate (x-axis) for the *tradeSeq* for the 100 cell trifurcating lineage simulation. The first panel displays overall performance, while the additional panels stratify genes into fifths based on quantification uncertainty as measured by the *InfRV* statistic averaged across cells. These results use 100 bootstrap replicates instead of 20.

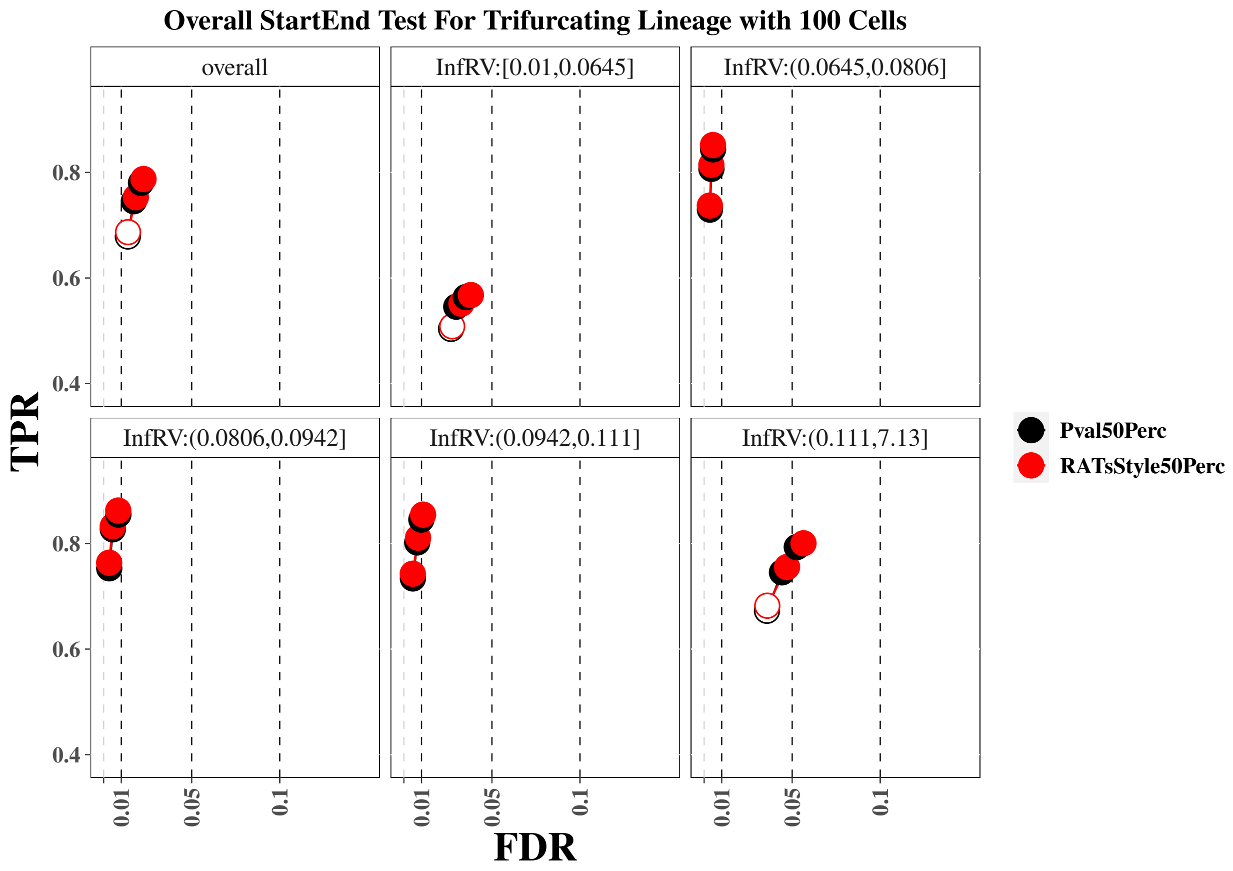


**Supplementary Figure S22:** True positive rate (y-axis) over false discovery rate (x-axis) for the *tradeSeq* for the 100 cell trifurcating lineage simulation. The first panel displays overall performance, while the additional panels stratify genes into fifths based on quantification uncertainty as measured by the *InfRV* statistic averaged across cells. These results compare our proposed Pval50Perc approach, which combines results across pseudo-replicates on the *raw* $p$-value scale, to an approach similar to that proposed in *RATs* that combines results across pseudo-replicates on the *adjusted* $p$-value scale ("RATsStyle50Perc"). See Section 2.4 in the main paper for more details about the differences between the two approaches.

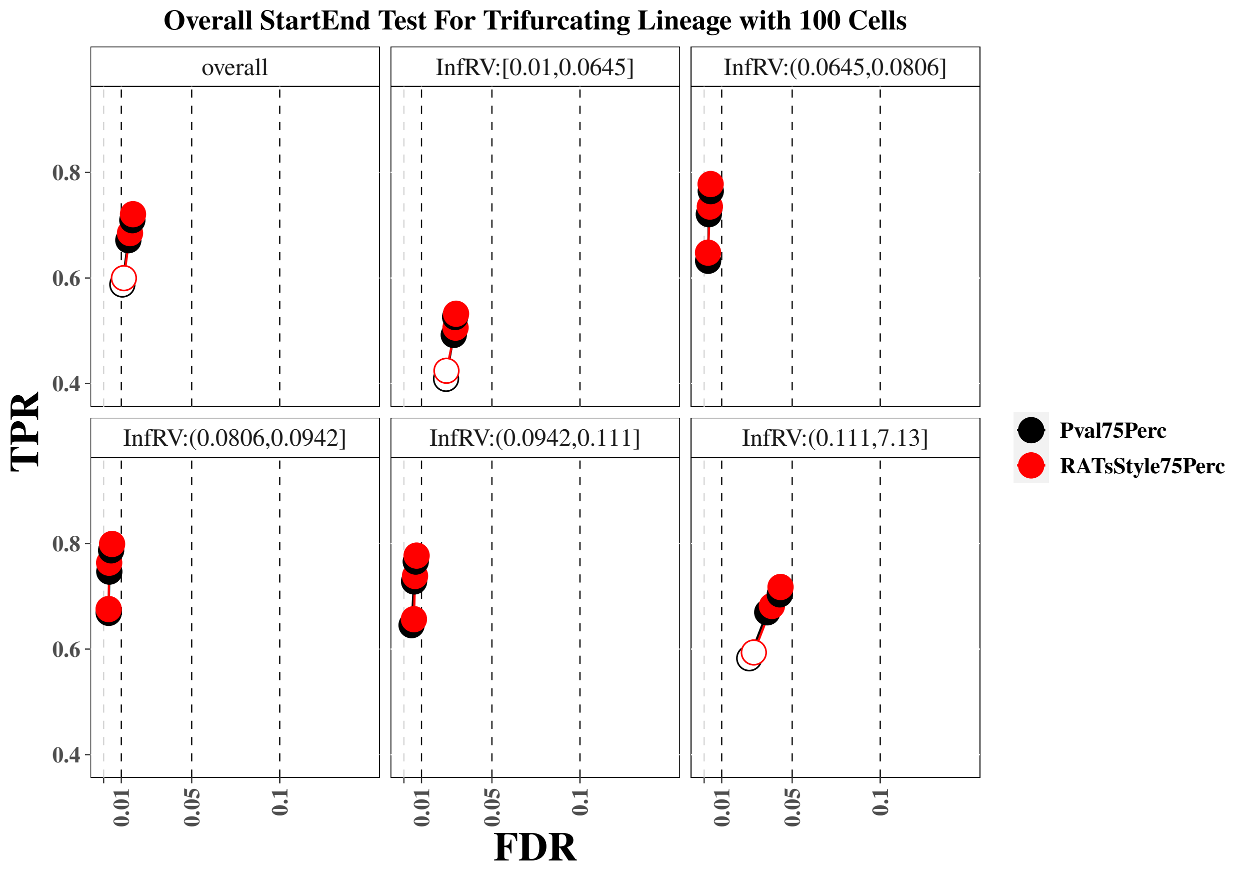


**Supplementary Figure S23:** True positive rate (y-axis) over false discovery rate (x-axis) for the *tradeSeq* for the 100 cell trifurcating lineage simulation. The first panel displays overall performance, while the additional panels stratify genes into fifths based on quantification uncertainty as measured by the *InfRV* statistic averaged across cells. These results compare our proposed Pval75Perc approach, which combines results across pseudo-replicates on the *raw* $p$-value scale, to an approach similar to that proposed in *RATs* that combines results across pseudo-replicates on the *adjusted* $p$-value scale ("RATsStyle75Perc"). See Section 2.4 in the main paper for more details about the differences between the two approaches.

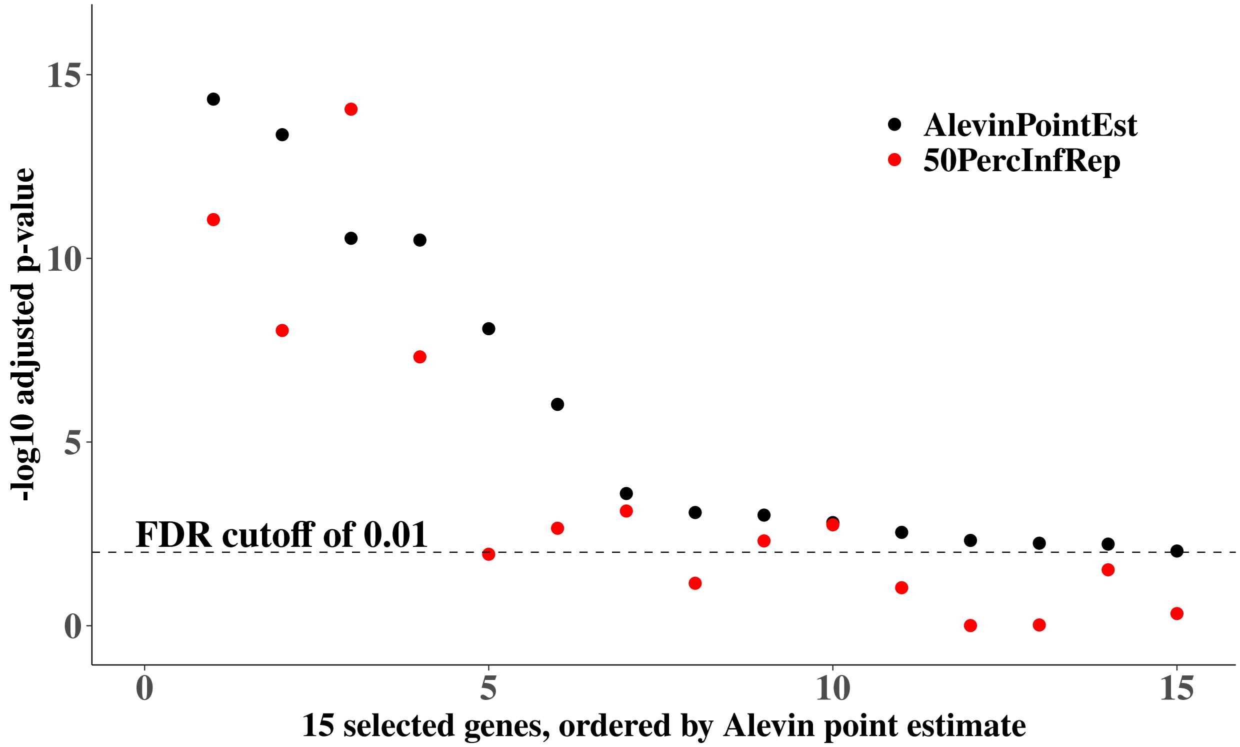


**Supplementary Figure S24:** Adjusted $p$-values for the `startVsEndTest` from *tradeSeq* for the trifurcating trajectory simulation with 100 cells for 15 null genes. $P$-values derived using the *Alevin* point estimate counts are shown in black (AlevinPointEst) and $p$-values corresponding to use of the 50th percentile $p$-value from repeating the analysis on each of 100 inferential replicates are shown in red (50PercInfRep). These genes were chosen for having a mean count $> 5$ across all 100 cells and a MeanInfRV value $> 0.50$. For 14 of 15 genes, use of the "50PercInfRep" approach resulted in a larger $p$-value (lower on the -log10 scale) than taking the $p$-value from the counts derived using the *Alevin* point estimate. Use of the "50PercInfRep" approach additionally eliminated 7 of the 15 false positives at an FDR level of 0.01.

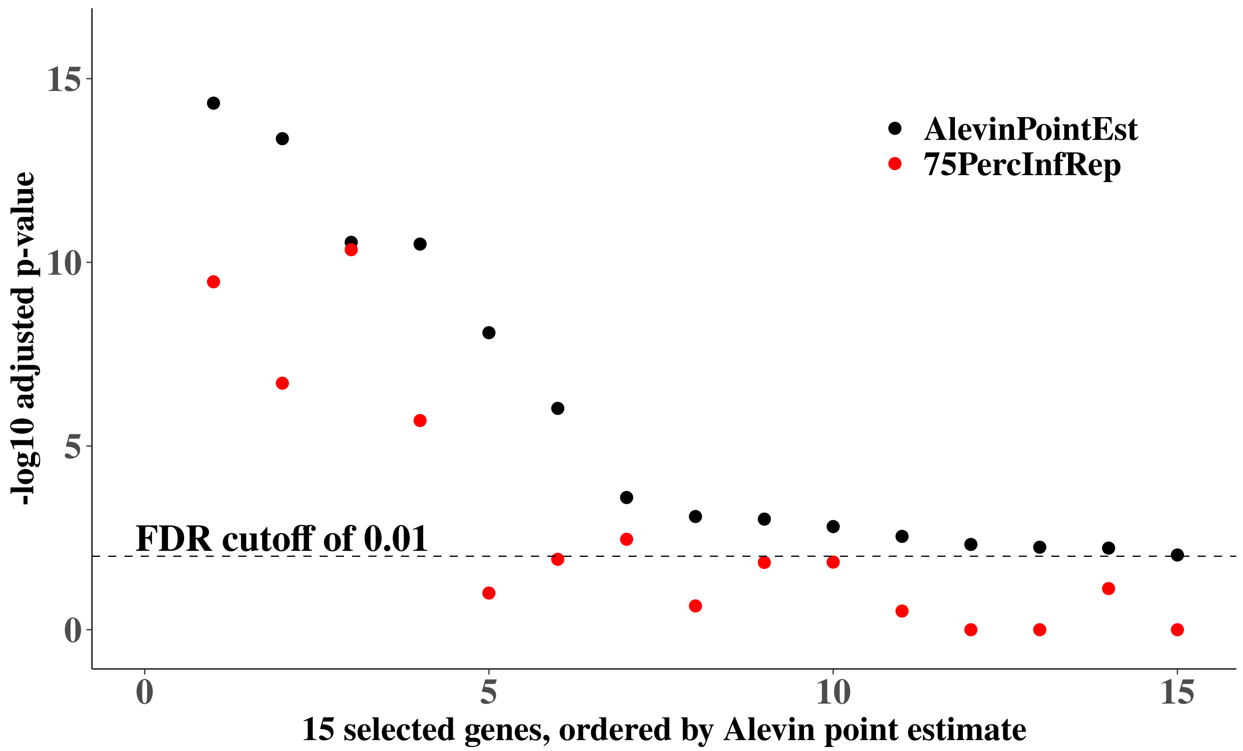


**Supplementary Figure S25:** Adjusted $p$-values for the `startVsEndTest` from *tradeSeq* for the trifurcating trajectory simulation with 100 cells for 15 null genes. $P$-values derived using the *Alevin* point estimate counts are shown in black (AlevinPointEst) and $p$-values corresponding to use of the 75th percentile $p$-value from repeating the analysis on each of 100 inferential replicates are shown in red (75PercInfRep). These genes were chosen for having a mean count $> 5$ across all 100 cells and a MeanInfRV value $> 0.50$. For all 15 genes, use of the "75PercInfRep" approach resulted in a larger $p$-value (lower on the -log10 scale) than taking the $p$-value from the counts derived using the *Alevin* point estimate. Use of the "75PercInfRep" approach additionally eliminated 10 of the 15 false positives at an FDR level of 0.01.

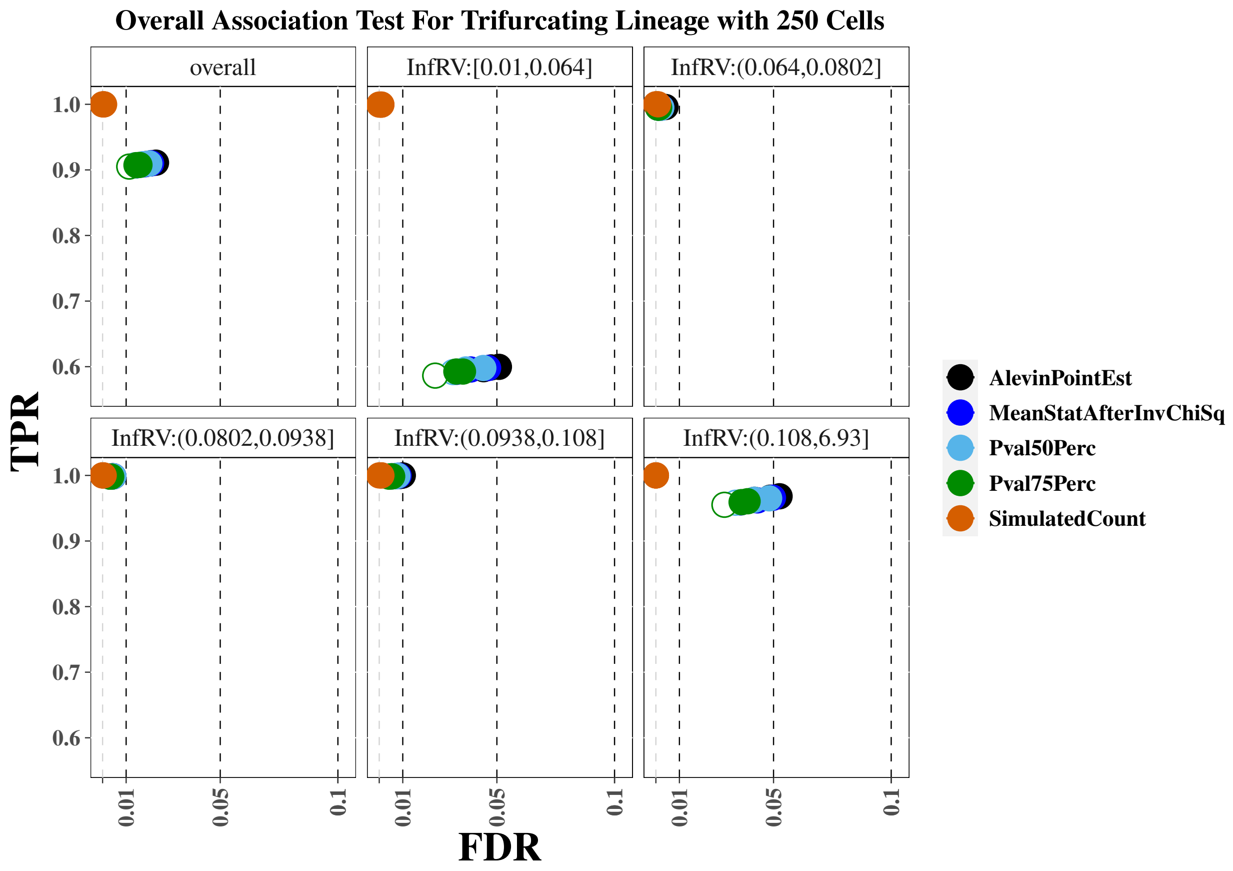


**Supplementary Figure S26:** True positive rate (y-axis) over false discovery rate (x-axis) for the *tradeSeq* for the 100 cell trifurcating lineage simulation. The first panel displays overall performance, while the additional panels stratify genes into fifths based on quantification uncertainty as measured by the *InfRV* statistic averaged across cells.

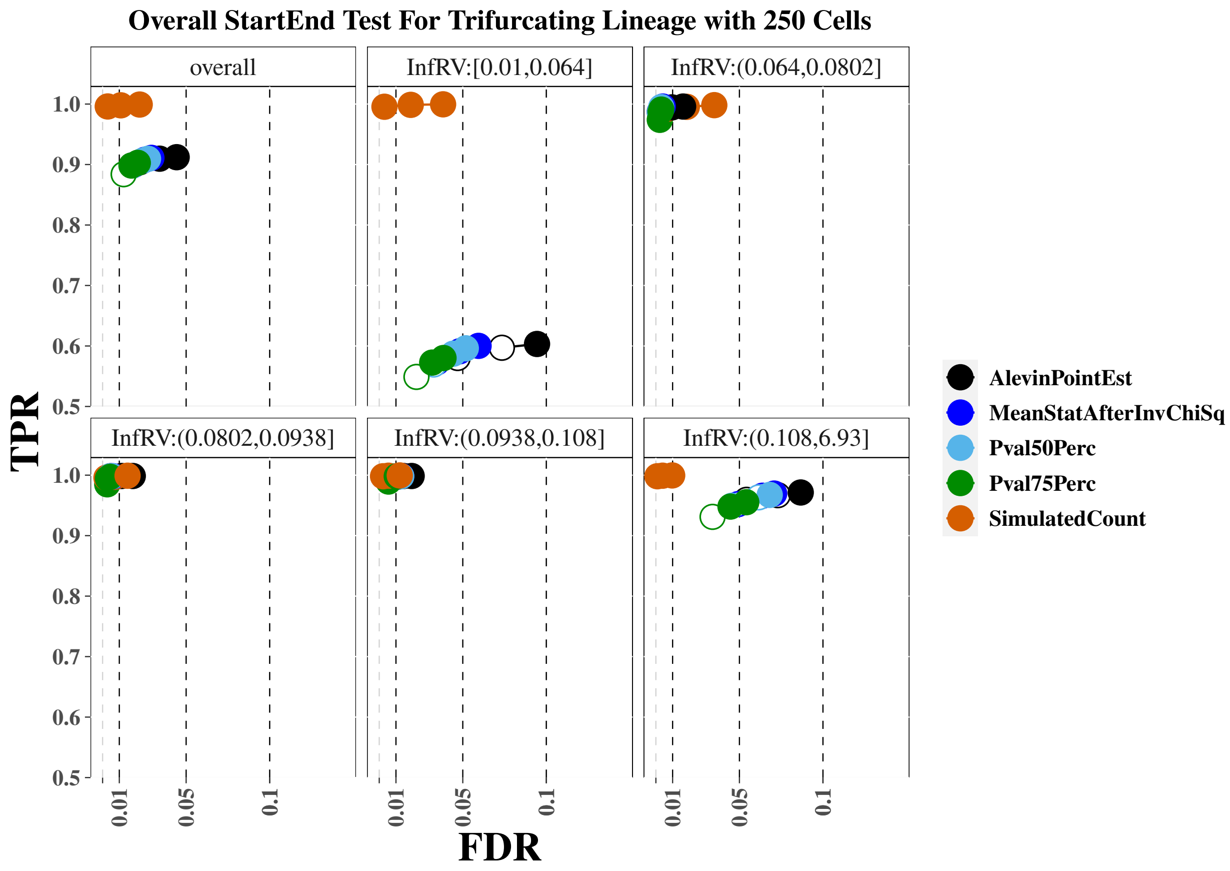


**Supplementary Figure S27:** True positive rate (y-axis) over false discovery rate (x-axis) for the *tradeSeq* for the 100 cell trifurcating lineage simulation. The first panel displays overall performance, while the additional panels stratify genes into fifths based on quantification uncertainty as measured by the *InfRV* statistic averaged across cells.

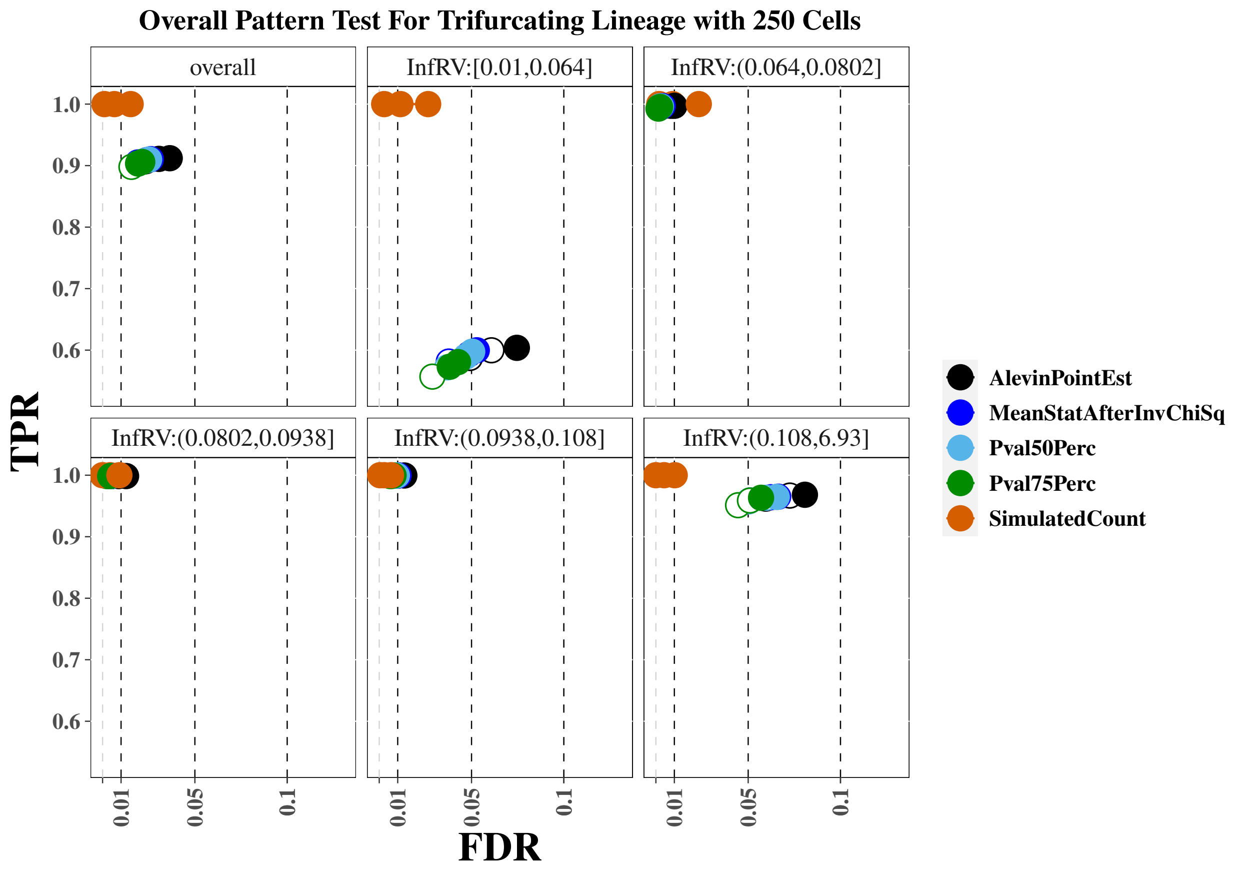


**Supplementary Figure S28:** True positive rate (y-axis) over false discovery rate (x-axis) for the *tradeSeq* for the 100 cell trifurcating lineage simulation. The first panel displays overall performance, while the additional panels stratify genes into fifths based on quantification uncertainty as measured by the *InfRV* statistic averaged across cells.

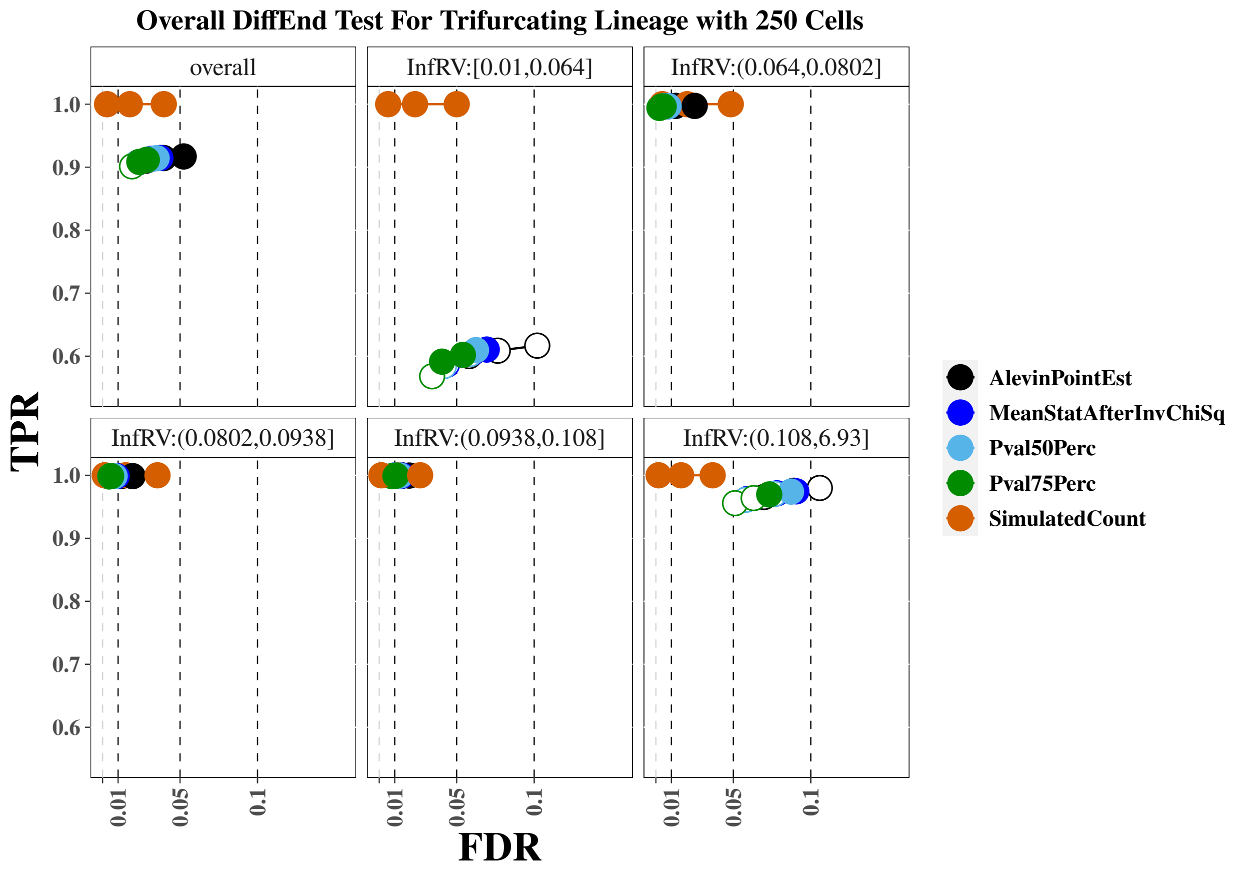


**Supplementary Figure S29:** True positive rate (y-axis) over false discovery rate (x-axis) for the *tradeSeq* for the 100 cell trifurcating lineage simulation. The first panel displays overall performance, while the additional panels stratify genes into fifths based on quantification uncertainty as measured by the *InfRV* statistic averaged across cells.

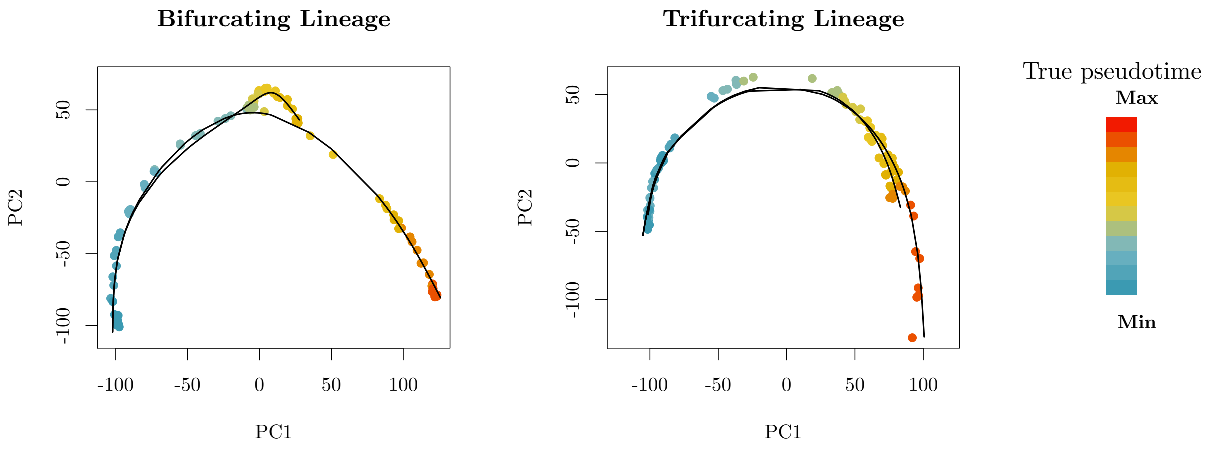


**Supplementary Figure S30:** Trajectory plots for the bifurcating and trifurcating lineages across the true pseudotime for the 100 cell simulations. The black lines plot the fitted lineages using *slingshot*. Code to plot this figure was taken from similar code used in *tradeSeq*.

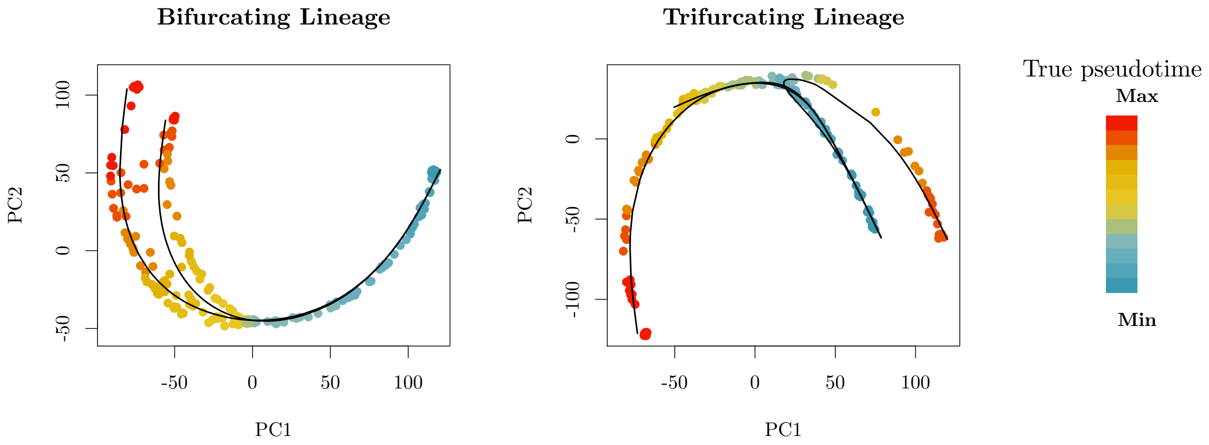


**Supplementary Figure S31:** Trajectory plots for the bifurcating and trifurcating lineages across the true pseudotime for the 250 cell simulations. The black lines plot the fitted lineages using *slingshot*. Code to plot this figure was taken from similar code used in *tradeSeq*.

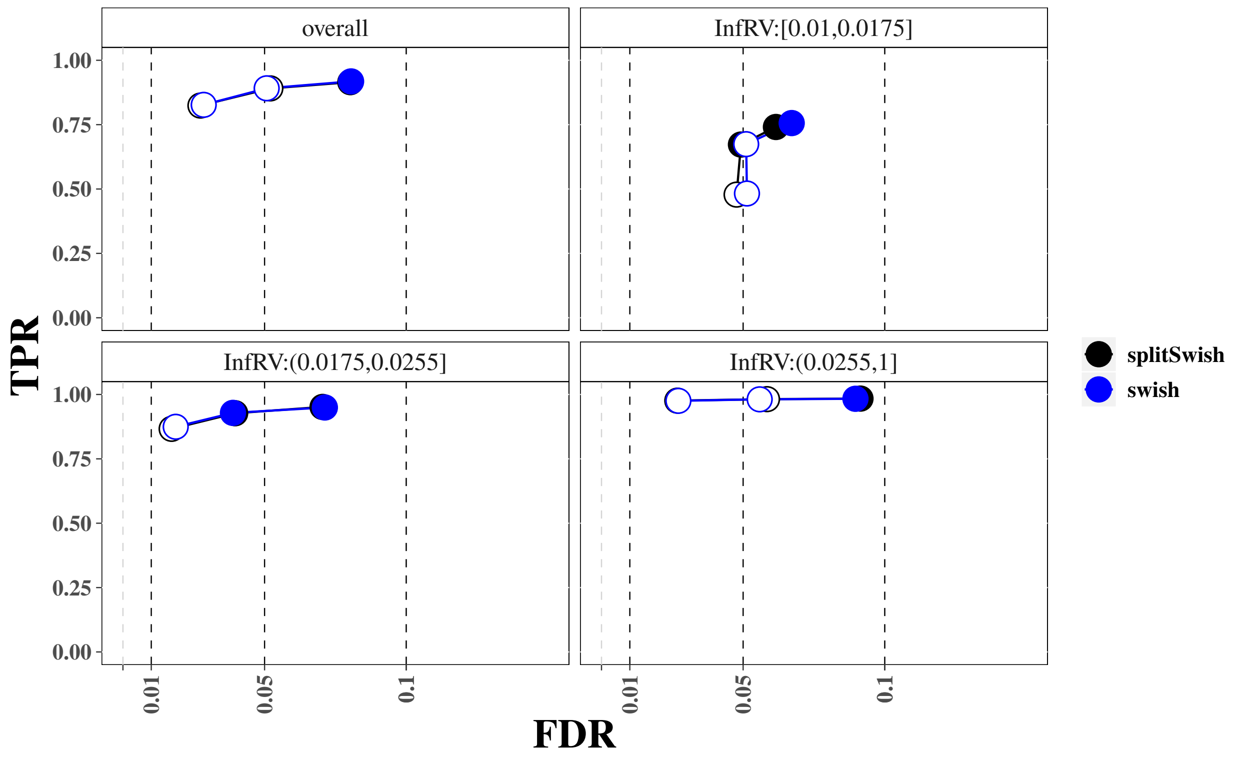


**Supplementary Figure S32:** Comparison of sensitivity and FDR for *swish* and *splitSwish* for the two group difference simulation.

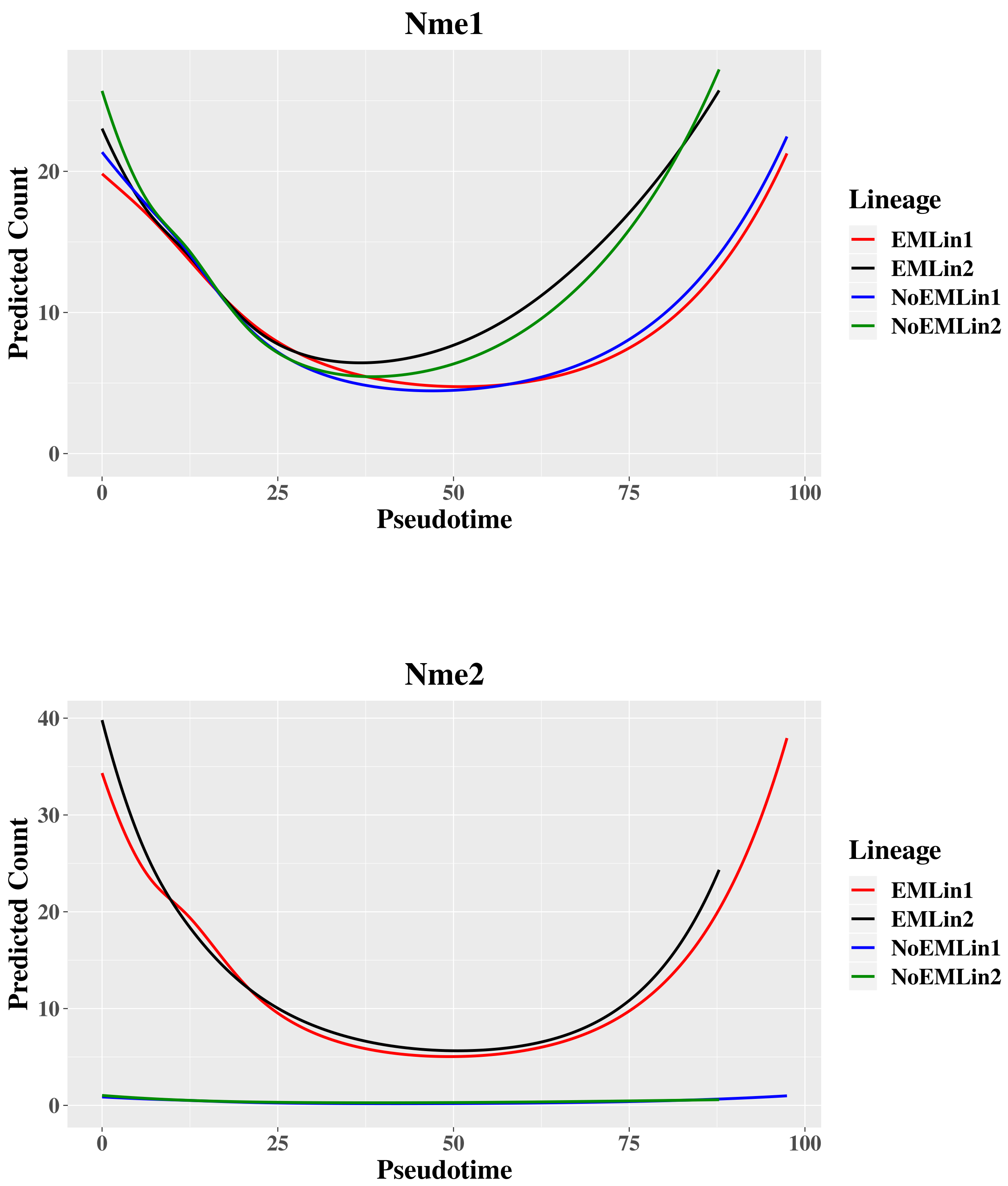


**Supplementary Figure S33:** Predicted counts over pseudotime for Nme1 and Nme2 for each lineage for results that incorporate multi-mapping reads via the EM algorithm and results that do not.

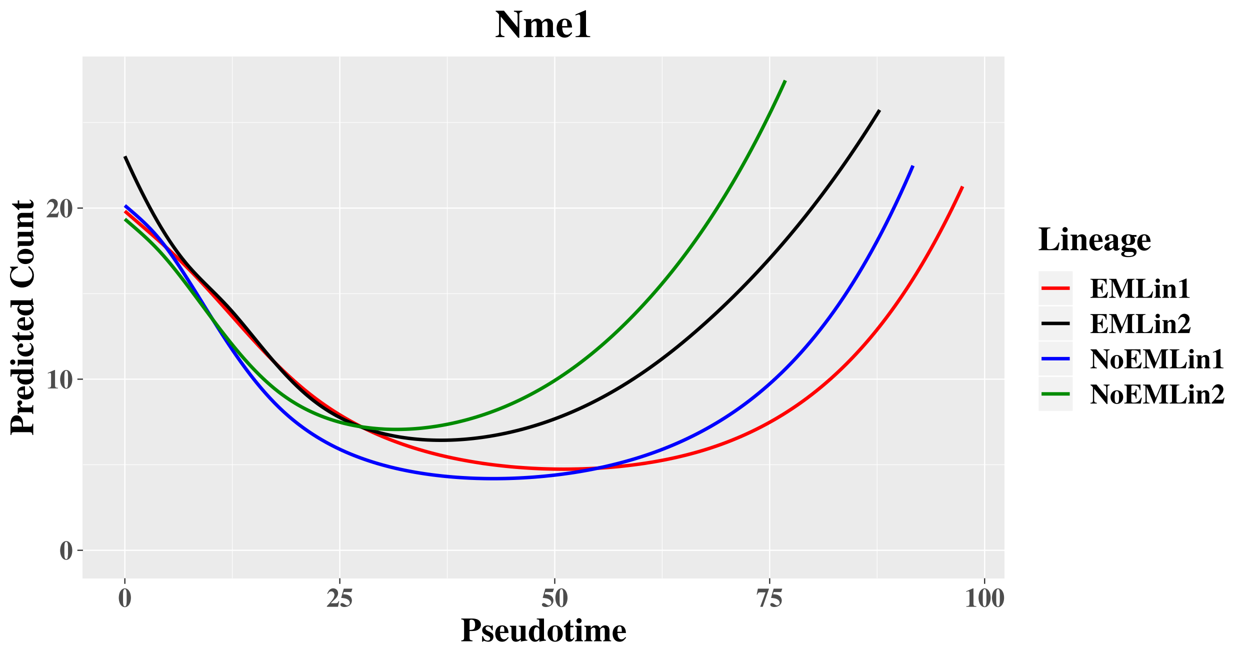


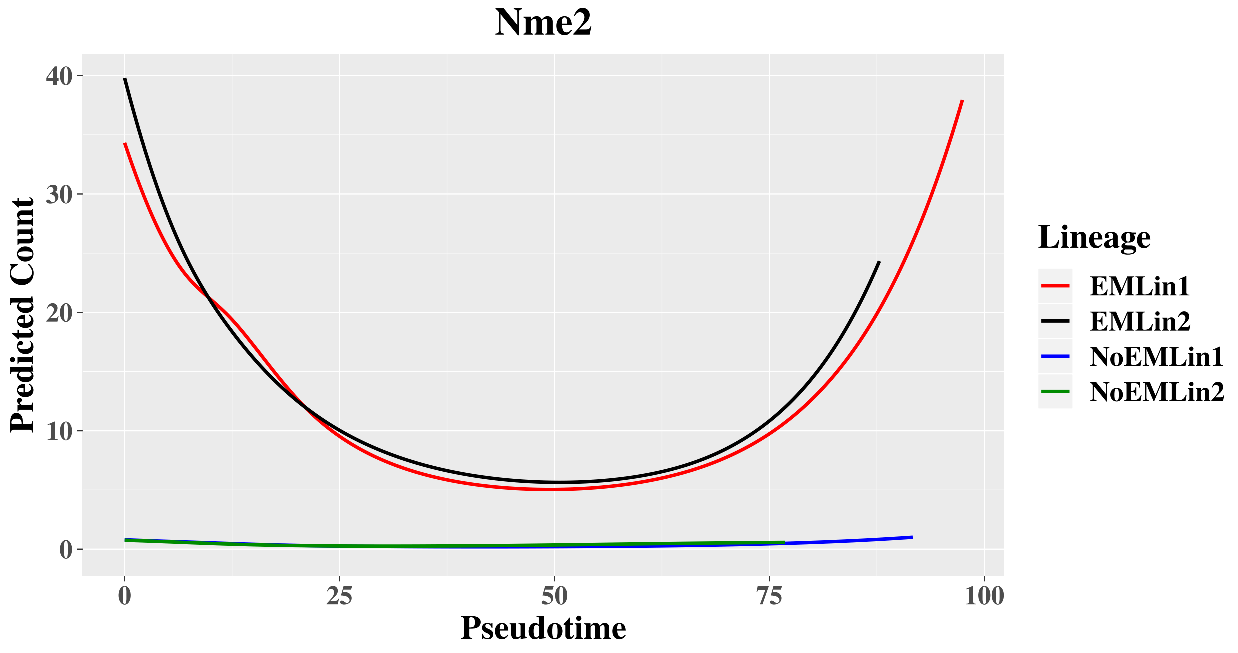


**Supplementary Figure S34:** Predicted counts over pseudotime for Nme1 and Nme2 for each lineage for results that incorporate multi-mapping reads via the EM algorithm and results that do not. Fit lineages in this case are allowed to differ between counts fit with and without the EM and EM algorithm, though the cell cluster assignments are fixed to be the same to ensure lineages “1” and “2” retain the same and are easily comparable to the results in the previous figure that force the lineages to be the same for the counts estimated with and without EM.

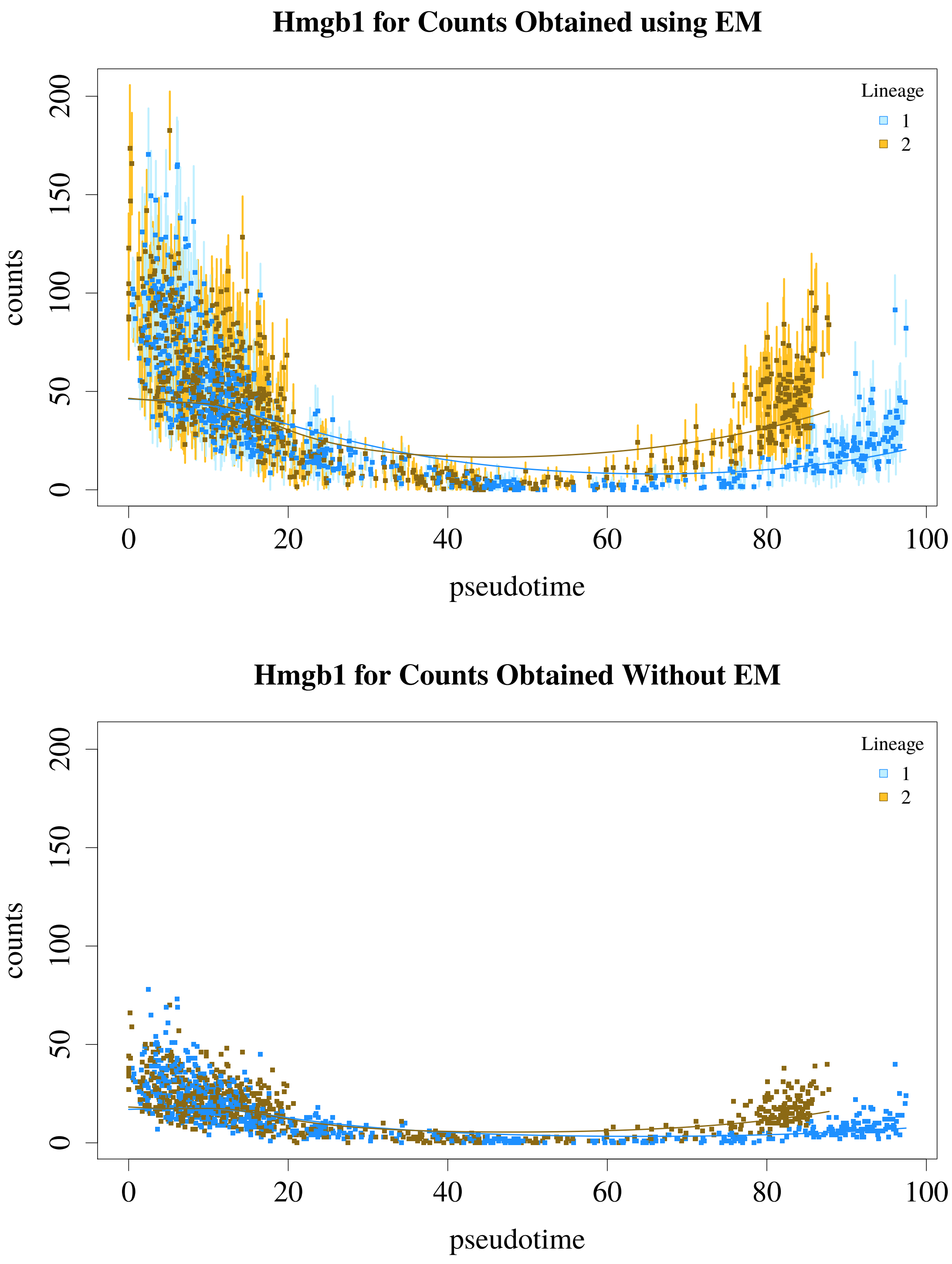


**Supplementary Figure S35:** Counts for Hmgb1 estimated incorporating multi-mapping reads with the EM algorithm (top panel) and without use of the EM algorithm (bottom panel). Both panels plot the fitted lineages for each cell, and the top panel additionally plots the uncertainty associated with each count using the bootstrap replicates.

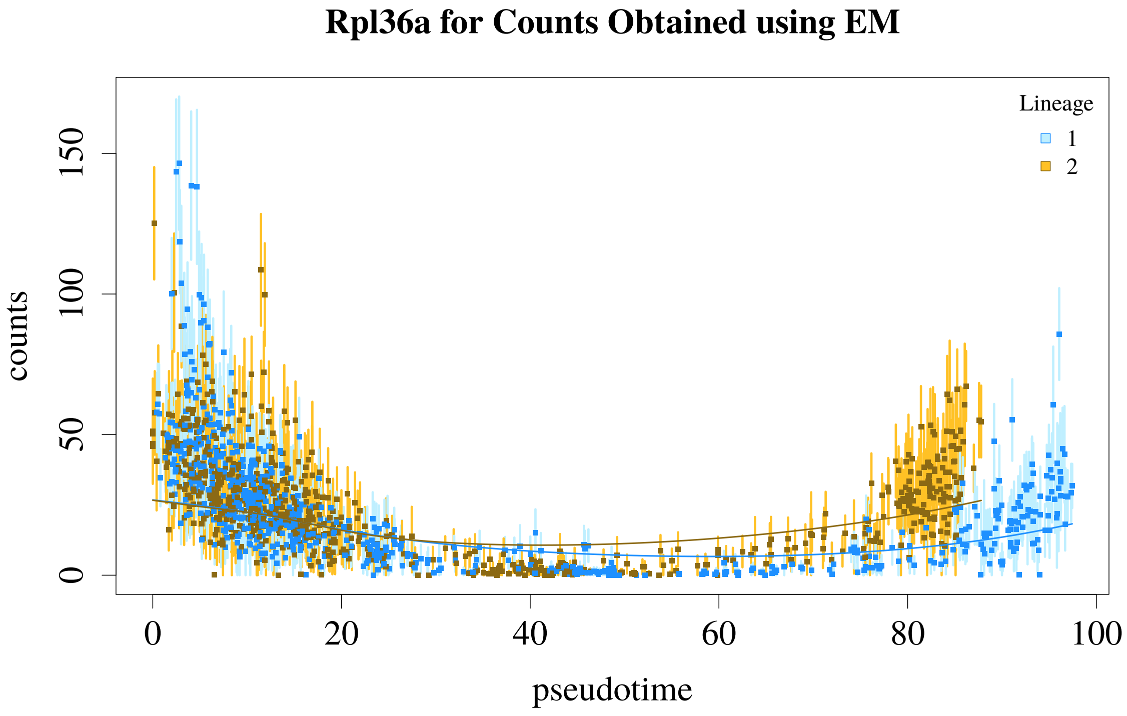


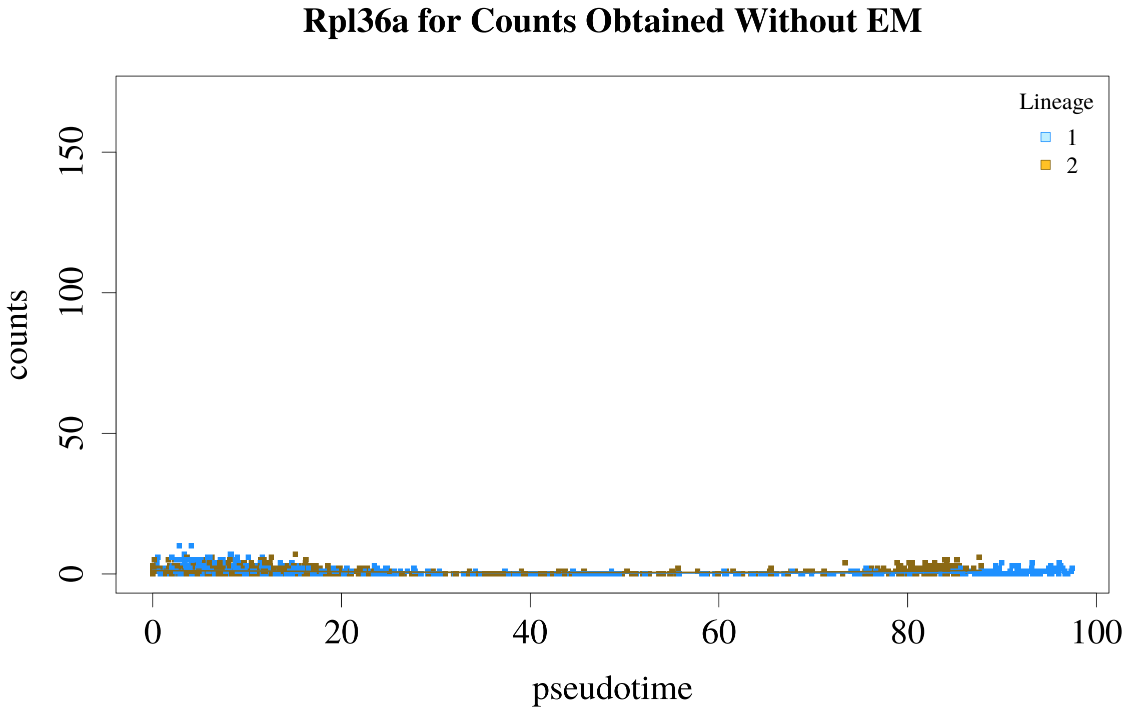


**Supplementary Figure S36:** Counts for Rpl36a estimated incorporating multi-mapping reads with the EM algorithm (top panel) and without use of the EM algorithm (bottom panel). Both panels plot the fitted lineages for each cell, and the top panel additionally plots the uncertainty associated with each count using the bootstrap replicates.

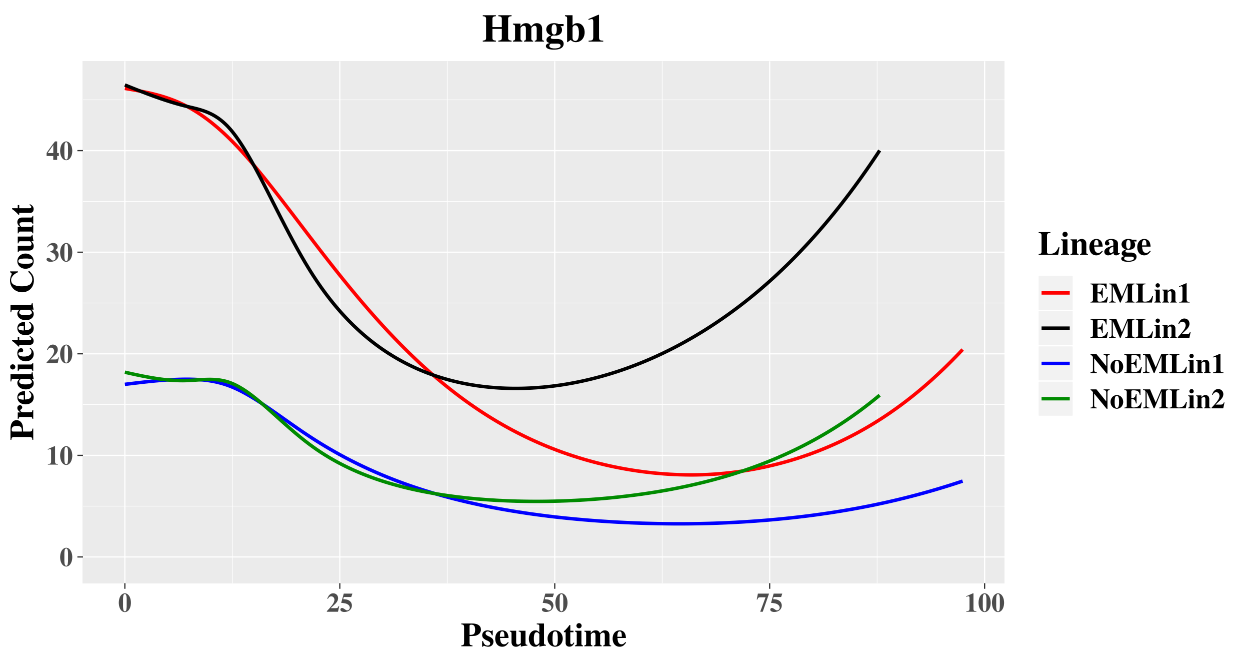


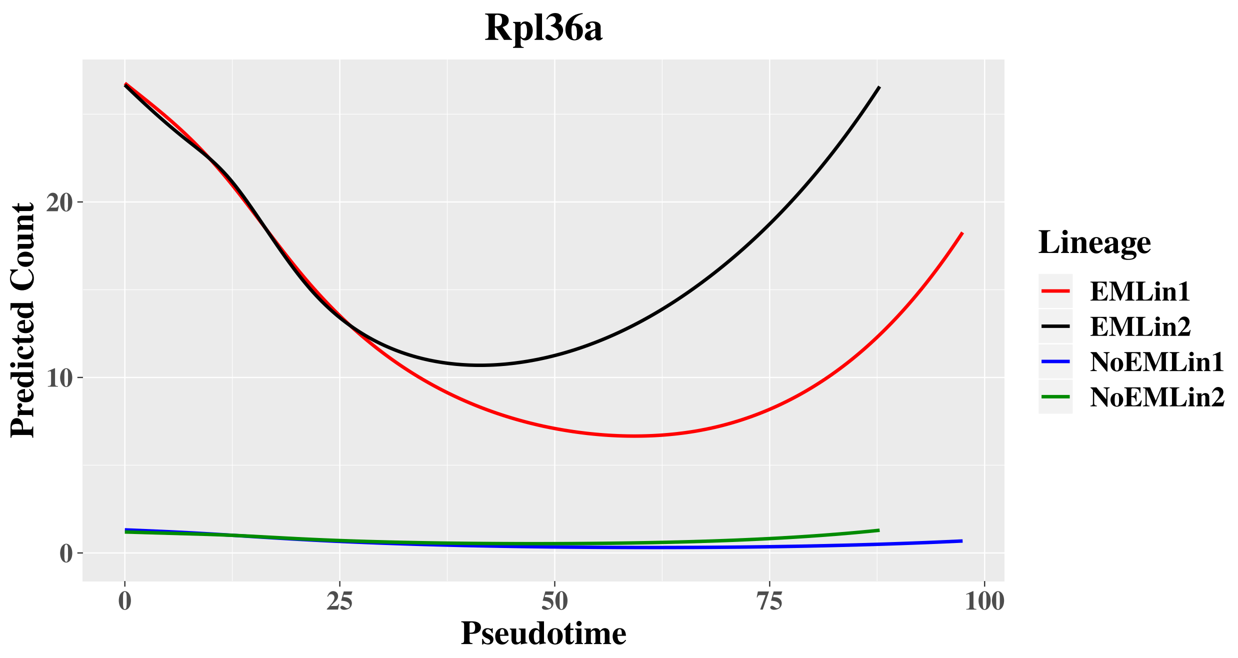


**Supplementary Figure S37:** Predicted counts over pseudotime for Hmgb1 and Rpl36a for each lineage for results that incorporate multi-mapping reads via the EM algorithm and results that do not.

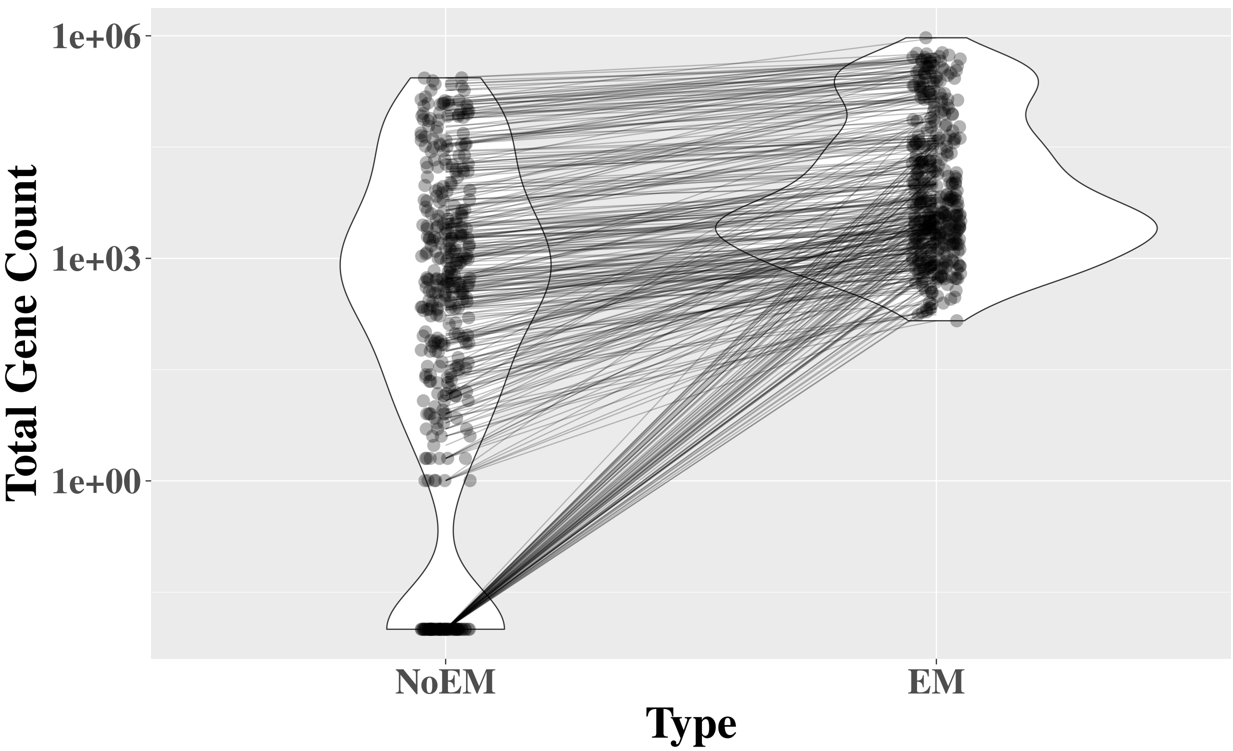


**Supplementary Figure S38:** Violin plot comparing counts from the 8.25 day time point of the mouse embryo data generated both incorporating multi-mapping reads ("EM") and not incorporating multi-mapping reads ("NoEM"). The y-axis is the total count of a particular gene summed across all 5,620 cells from the 8.25 day time point. The 358 genes plotted have a count of at least 3 in at least 10 cells and have a $\log_2$ ratio between the EM and NoEM counts that is larger in absolute value than $\log_2(1.5)$. Values for the same gene from the NoEM and EM types are connected.
